# High-throughput viscoelastic characterization of cells in hyperbolic microchannels

**DOI:** 10.1101/2023.12.10.571005

**Authors:** Felix Reichel, Ruchi Goswami, Salvatore Girardo, Jochen Guck

## Abstract

Extensive research has demonstrated the potential of cell viscoelastic properties as intrinsic indicators of cell state, functionality, and disease. For this, several microfluidic techniques have been developed to measure cell viscoelasticity with high-throughput. However, current microchannel designs introduce complex stress distributions on cells, leading to inaccuracies in determining the stress-strain relationship and, consequently, the viscoelastic properties. Here, we introduce a novel approach using hyperbolic microchannels that enable precise measurements under a constant extensional stress and offer a straightforward stress-strain relationship, while operating at a measurement rate of up to 100 cells per second. We quantified the stresses acting in the channels using mechanical calibration particles made from polyacrylamide (PAAm) and found that the measurement buffer, a solution of methyl cellulose and phosphate buffered saline, has a constant extensional viscosity of 0.5 Pa s up to 200 s^-1^. By measuring oil droplets with varying viscosities, we successfully detected changes in the relaxation time of the droplets and our approach could be used to get the interfacial tension and viscosity of liquid-liquid droplet systems from the same measurement. We further applied this methodology to PAAm microgel beads, demonstrating the accurate recovery of Young’s moduli and the near-ideal elastic behavior of the beads. To explore the influence of altered cell viscoelasticity, we treated HL60 human leukemia cells with Latrunculin B and Nocodazole, resulting in clear changes in cell stiffness while relaxation times were only minimally affected. In conclusion, our approach offers a streamlined and time-efficient solution for assessing the viscoelastic properties of large cell populations and other microscale soft particles.

## Introduction

Cellular viscoelasticity has emerged as a central indicator of cell state and function, playing an increasingly significant role in disease characterization.^1–6^ Established techniques such as micro-pipette aspiration,^1,7^ atomic force microscopy,^8–10^ or the optical stretcher,^11–14^ while effective in revealing changes in cell deformability and relaxation times, operate on timescales of seconds with throughputs limited to a few hundred cells per hour max. This limitation makes them unsuitable for swift diagnostic applications.

To address the need for faster measurements, microfluidic techniques have gained prominence.^3,5,15–17^ Operating at timescales ranging from below 1 to 100 milliseconds, these platforms offer measurement rates of over 1,000 cells per second. However, the quantification of stresses and cell strain in the microchannels remains challenging due to the complex three-dimensional stress distribution on the cell surface.^18,19^

Viscoelastic analysis typically necessitates a one-dimensional stress-strain relationship to derive mechanistic properties such as elastic moduli, relaxation times, or storage and loss moduli.^5,10,20,21^ Moreover, in cell mechanics measurements, polymeric crowding agents are often employed to enhance the viscosity of the cell carrier solution.^5,16,22,23^ This augmentation leads to non-Newtonian behavior, including shear-thinning, the occurrence of normal stress differences, or strain thickening.^24–26^ Notably, this behavior is frequently characterized solely in shear, neglecting the qualitatively different behavior in extension. ^5,22,23^

In this study, we address these challenges by introducing hyperbolic microchannels as a novel approach for accurate cell viscoelastic measurements under well-defined tensile stresses and strains. The flow pattern within the hyperbolic channel geometry creates a region of constant stress, simplifying the analysis of experiments. While hyperbolic channels have been used before for cell mechanical measurements, the focus has predominantly been on measuring cell deformability, with little attention given to correlating stresses to strains within the hyperbolic region to quantify cell viscoelastic properties.^22,27,28^

Our study aims to fill this gap by simultaneously examining both cell deformability and associated stresses within hyperbolic microchannels. Additionally, our methodology extends to the determination of extensional stresses in polymeric solutions at varying extension rates.

To underscore the method’s capability in measuring changes in relaxation times, we conducted experiments using silicone oil droplets of varying viscosities. Our results not only showcase the ability to detect changes in timescales but also highlight the method’s utility in retrieving interfacial tension and droplet viscosity from a single experiment.

Employing a Kelvin-Voigt model, we analyze the stress-strain relationship to extract the Young’s modulus and relaxation times of cells. Validation of our approach involves measurements on polyacrylamide (PAAm) microgel beads,^29^ demonstrating the recovery of expected Young’s moduli and indicating near-elastic behavior based on relaxation times.

Finally, we use the hyperbolic microchannels to investigate changes in HL60 cell viscoelasticity under treatment with the actin polymerization inhibiting drug Latrunculin B (LatB) and microtubule depolymerizing reagent Nocodazole. Our findings reveal a concentration-dependent decrease in the cells’ Young’s modulus, while relaxation times exhibit only slight alterations. In conclusion, our results underscore the utility of hyperbolic microchannels for accurate measurements of cell viscoelastic properties in extension at up to 100 cells per second with potential applications in cell research and diagnostics.

## Theoretical background

### Stresses and deformations in extensional flow

During flow through a converging channel in *x*-direction with a constant flow rate, *Q*, the fluid velocity, *v*, will increase over *x*. The rate of this velocity increase is called the *extension rate* 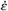:

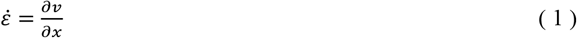

A planar, extensional flow field is defined with the following velocity components:

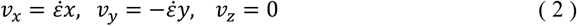

at a constant extension rate. This results in a tensile stress component in *x*-direction, *σ*_*x*_, and compressive stress component in *y*-direction, *σ*_*y*_ (see Fig. 1A). The net stress acting on an object in this flow field is defined as:

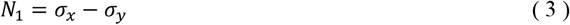

The measure *N*_1_ is known in rheology as the *first normal stress difference* and is a material property of a liquid.^30,31^ The stress in *z*-direction is zero: *σ*_*z*_ = 0. For Newtonian liquids, there is a linear relationship between *N*_1_, the extension rate, and the shear viscosity, η. For planar extension, the case defined by equation (2), the relation reads 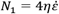. For non-Newtonian liquids, *N*_1_ usually cannot be derived from *η* and the behavior in extension has to be characterized independently. Generally, the relation between *N*_1_ and the extension rate is described by the *extensional viscosity* 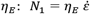 and *η*_*E*_ will often be dependent on 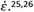.^25,26^

**Figure 1:**
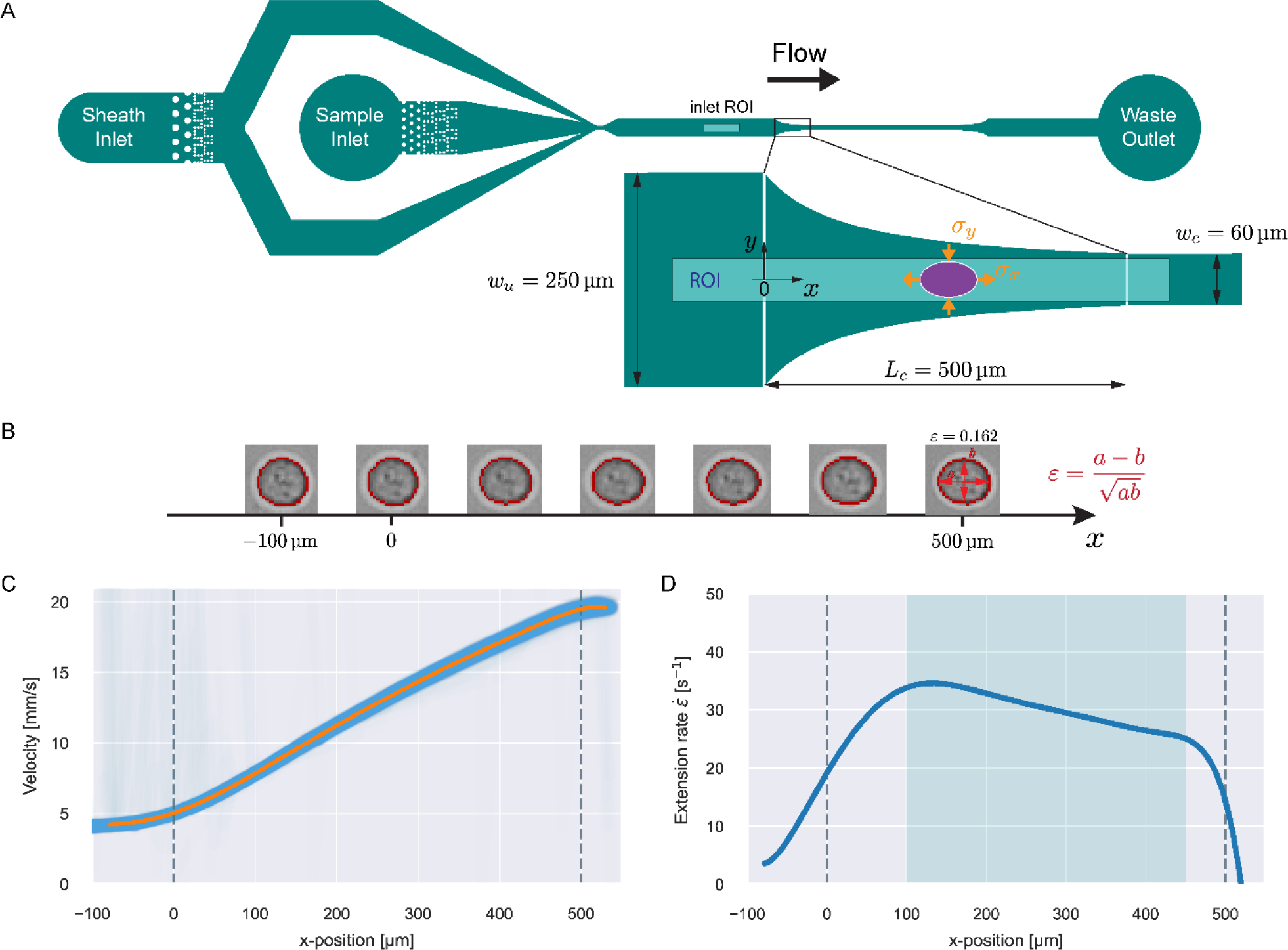
Microfluidic channel design and measurement principle. **A**) Full design of the microfluidic channel with magnified illustration of the hyperbolic region. The ellipse represents a deformed particle and the stresses in *x* and *y* are depicted. The shaded region indicates the ROI for a viscoelasticity measurement in the hyperbolic region. The inlet ROI shown in the full design indicates the region for inlet measurements to correct for optical distortions. **B**) Trajectory of an exemplary HL60 cell flowing through the hyperbolic region. The detected contours are shown in red. At the contour of the right-most cell, the strain calculation is explained. **C**) Velocity trajectories as function of *x* for single 370 Pa PAAm beads (blue lines) at 0.02 μL/s flow rate. The orange line shows a polynomial fit to the data. **D**) Extension rate of the fitted velocity curve in C as function of *x*. The shaded region highlights the zone of consistent extension rate.

For this stress definition, we can define a corresponding net strain as:

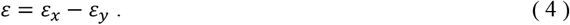

Assuming that the particle is initially a sphere, it is expected to deform into an ellipsoidal shape, in first order approximation, when flowing in the extensional field.^32–35^

We define the strain in *x, y* by the deviation of the contour from the initial sphere radius *R*_0_:

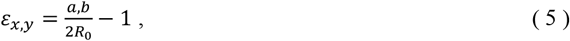

where *a, b* are the major and minor axes of the ellipse. The strain in *z*-direction is expected to be zero: *ε*_*z*_ = 0. The resulting net strain can be computed from *a* and *b*, assuming the volume of the particle stays constant during the deformation:

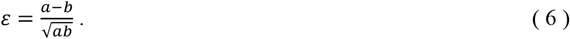

The shapes and strain of an example HL60 cell flowing through the extensional field is shown in figure 1B.

### Deformation of liquid droplets in extensional flow

The deformation of a liquid droplet flowing in an extensional field with interfacial tension, *γ*, droplet viscosity, *η*_drop_, and viscosity of the carrier solution, *η*_0_, is well described in the literature.^32,33,36^ Here, we use the derivations from Cabral and Hudson to compute *γ* and *η*_drop_ from the droplet deformation and extension rate.^36^

The droplet deformation is defined as the Taylor deformation:^32^

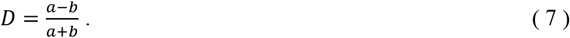

The interfacial tension at the steady-state deformation, *D*_*∞*_, can be computed with:

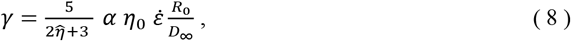

with the viscosity ratio 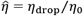, and the effective viscosity *α*:

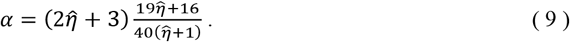

In the absence of flow or at uniform flow velocity, a droplet with deformation *D*_0_ at time *t* = 0 will relax towards equilibrium following an exponential:^36^

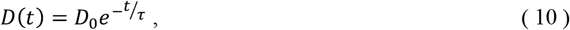

with the droplet relaxation time, τ. The relaxation time is related to the other system parameters via:^33,36^

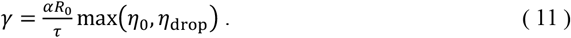

τ and *D*_*∞*_ can be derived from the time evolution of the droplet deformation at constant extension rate. We assume that *R*_0_ and *η*_0_ are known and that *η*_0_ < *η*_drop_. *γ* and *η*_*drop*_ can be derived by equating equations 8 and 11 (see SI text).

### Kelvin-Voigt model analysis

We characterized the viscoelasticity of microgel beads and cells using the Kelvin-Voigt material model:

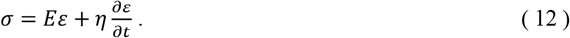

Here, *σ* is a one-dimensional stress, *E* an apparent elastic modulus and *η* an apparent bulk viscosity. For a general stress function in time, *σ*(*t*), and an initial strain, *ε*_0_, at *t* = 0, the general solution of the model is:

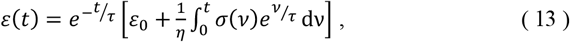

with the relaxation time τ = *η*⁄*E*.

## Materials and Methods

### Microfluidic chip design

The entire chip geometry is shown in figure 1A. Flow is introduced via FEP-tubing into the sheath and sample inlet. The sample particles are only delivered to the channel through the sample inlet and the sheath only contains the carrier solution. At the point where sheath and sample flow meet, the sample particles get focused into the channel center before the measurement region (shown as zoomed-in inset). The dotted structures after the inlets show filter pillars that prevent large particles or particle aggregates from clogging or contaminating the measurement region.

After sheath focusing, the channel continues straight for 2.4 cm. This was introduced to give particles enough time to relax from the deformation that is induced by the sheath focusing before reaching the measurement region. Channel height of all chips used for this study was 30 μm. This height was chosen to ensure cells with an approximate diameter of 15 μm will be focused at the flow centerline. For higher channels, cells would take up a lateral equilibrium position off the centerline.^37,38^

The measurement region, also called hyperbolic region, was designed to ensure a linear increase of the centerline velocity along the *x* -axis. The centerline velocity, *u*_0_, was computed with the following equation:^39^

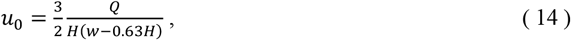

with flow rate *Q*, channel height, *H*, and channel width, *w*, where *w* = *w*(*x*). Equation 14 is a good approximation for *u*_0_ when *w* ≫ *H*. To ensure a constant extension rate 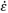, *w*(*x*) must be constructed such that 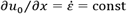. This leads to the following equation for the channel width in the hyperbolic region:

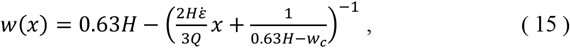

with the contraction width, *w*_*c*_ (see Fig 1A and SI text).

A channel geometry is fully defined by the upper contraction width, *w*_*u*_, lower contraction width, *w*_*c*_, and contraction length, *L*_*c*_. The measures relate to the flow parameters with the following relation:

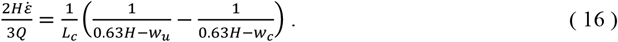

The resulting velocity as function of *x* for PAAm microgel beads with a Young’s modulus of 370 Pa at a flow rate of 0.02 μL/s are shown in figure 1C. The blue lines depict velocity trajectories of single beads and the orange line shows a 12^th^-order polynomial fit to the data. The extension rate for the sample was computed from the fitted velocity curve and is shown in figure 1D.

After the hyperbolic contraction, the channel continues straight for about 2 cm to observe the relaxation from the previous stress. Before the waste outlet, an expansion region mirroring the hyperbolic contraction was added to observe objects in expansion.

### Device fabrication

Microfluidic chips were produced using standard soft lithography methods with polydimethylsiloxane (PDMS). To produce the master mold, a 30-μm-thick layer of AZ® 15nXT (450 CPS) photoresist (MicroChemicals GmbH, Germany) was spin-coated onto a 4-inch silicon wafer (2000 rpm, 5000 rpm/s, for 2.8 s) using a spin coater (WS-650Mz-23NPP, Laurell, USA) and soft-baked at 110 °C for 3 minutes. The chip design, depicted on a chromium photomask, was transferred on the coated substrate via UV exposure at 600 mJ/cm^2^ (MA6 Gen4, Suess MicroTec GmbH, Germany), followed by a post-baking at 120 °C for 1 minute and development in a 1:3 dilute solution of AZ® 400 K developer (MicroChemicals GmbH, Germany) in deionized water for 150 seconds. The final master mold received a hydrophobic coating through vapor deposition of 1H,1H,2H,2H-perfluorooctyl-trichlorosilane (Sigma-Aldrich 448931, CAS 78560-45-9) in a vacuum desiccator for 1 day, facilitating the easy peel-off of replicas during replica molding. PDMS (base to curing agent ratio of 10:1 w/w; VWR 634165S, SYLGARD 184) was poured over the master, degassed, and cured at 78 °C for 1 hour and 15 minutes. Following the cutting of the chips and the creation of 1.5 mm inlet and outlet holes, the PDMS replica was bonded to a glass cover slip (40×24 mm^2^, thickness 2, Hecht, Germany) through an air plasma treatment (Plasma system Atto, Diener Electronic, Germany) at 75 W for 2 min, 20 sccm air flow and 0.4 mbar pressure.

### Production of carrier medium

Measurements on oil droplets were performed by dispersing the concentrated droplet solution into a mixture of 83.8 v/v% glycerol (Sigma-Aldrich G5516, CAS 56-81-5) with water. This concentration of glycerol corresponds to a dynamic viscosity of 100 mPa s at 25 °C.^40,41^

Before measurements, cells and beads were suspended into a 0.6 w/w% solution of methyl cellulose (MC) (4000 cPs, Alfa Aesar 036718.22, CAS 9004-67-5) dissolved in phosphate buffered saline (PBS) (Gibco Dulbecco 14190144) following the protocol of Büyükurganci et al.^16,24^ The osmolality of PBS was adjusted to values between 280–290 mOsm/kg before the addition of MC (VAPRO Vapor Pressure Osmometer 5600, Wescor, USA). To prepare MC-PBS solutions for 1 L of PBS (1010 g), 6.00 g MC powder were utilized, resulting in a 0.594 w/w% MC concentration in the final solution. Following the addition of the powder, the mixtures were maintained at 4 °C under continuous rotation for two days to allow for the dissolution of MC. This process involved placing a rotating mixer (RS-TR05, Phoenix Instrument) in a High-Performance Pharmacy Refrigerator (TSX Series, Thermo Fisher Scientific), with the bottles containing the MC-PBS solution rotating inside. Subsequently, the pH value was adjusted to 7.4 (Orion Star A211 benchtop pH meter, Thermo Scientific) by the addition of NaOH. In the next step, the solutions were sterile-filtered using membrane filters (Merck Millipore Steritop-GP polyethersulfone (PES) membrane, 0.22 mm pore size). The buffer solutions were then stored at 4 °C.

### Latrunculin B and Nocodazole treatments

A stock solution of Latrunculin B (Sigma-Aldrich L5288, CAS 76343-94-7) was prepared by dissolving the powder in DMSO at a concentration of 10 mM. This stock solution was then further diluted in DMSO to 10,000 times the desired concentration, ensuring an equivalent DMSO concentration in all treatments (0.01 v/v%). Subsequently, LatB was diluted by a factor of 10,000 in a 0.6 w% MC-PBS buffer, resulting in final LatB concentrations of 0.1, 1, 5, 10, 25, 50, 100, and 250 nM. The same approach was used for Nocodazole (Sigma-Aldrich SML1665, CAS 31430-18-9) to get a final concentration of 1 μM.

### Production of PAAm-microgel beads

PAAm beads were produced according to the protocol introduced by Girardo et al.^29^ Briefly, a PDMS-based flow-focusing microfluidic chip was used to produce polyacrylamide pre-gel droplets in fluorinated oil (3M™ Novec™ 7500, Iolitec Ionic Liquids Technologies GmbH, Germany). The fluorinated oil contained ammonium Krytox® surfactant (2.4 w/v%) as the emulsion stabilizer and 0.4 v/v% N, N, N′, N′-tetramethylethylenediamine (TEMED) (Sigma-Aldrich T9281, CAS 110-18-9) as the catalyst. The pre-gel mixture contained acrylamide (40 w/v %) (Sigma-Aldrich A8887) as the monomer, bis-acrylamide (2 w/v%) (Sigma-Aldrich 146072) as the crosslinker, and ammonium persulphate (0.05 w/v%) (Cytiva GE17-1311-01) as the free radical initiator, diluted in 10mM Tris buffer (pH = 7.48). Total monomer concentration (%*c*_*T*_) in the pre-gel mixture together with the droplet diameter was fine-tuned to obtain beads with precise diameter and elasticity within the desired range. After in-drop polymerization at 65 °C, the beads were washed and resuspended in 1×PBS (pH = 7.4, Gibco), and ultimately stored at 4 °C.

A list of the PAAm beads used in this study is reported with both their total monomer concentration (%*cc*_*TT*_), mean diameter and Young’s modulus (Table S1). The Young’s modulus of the beads was measured with real-time deformability cytometry (RT-DC, see SI text).^16,19,24^

### Production of silicone oil droplets

The same chip design and setup employed in the PAAm bead production was also used for generating silicone oil droplets. This process involved a PDMS-based chip with a channel height of 14 μm and channel width of 15 μm at the cross junction. Before starting the droplet production, the microfluidic chips underwent a treatment by an air plasma (Plasma system Atto, Diener Electronic, Germany) at 75 W for 4 min, 20 sccm air flow and 0.4 mbar pressure. This treatment rendered the surface of the PDMS channel walls hydrophilic, a crucial step to enable on-chip production of an oil-in-water microemulsion.^42^

Two distinct types of droplets were generated using two silicone oils with kinematic viscosities of 500 and 1,000 cSt (Sigma-Aldrich 37380 & 378399, CAS 63148-62-9) as the dispersed phase. The continuous phase was a aqueous solution of 2 v/v% Poly(ethylene glycol) monooleate (Sigma-Aldrich 460176, CAS 9004-96-0) used as a emulsifier.

Droplets with a kinematic viscosity of 500 cSt were produced under a set pressure of 480 mbar for the continuous phase and 310 mbar for the dispersed phase, resulting in droplets with an average diameter of 16 μm at a production rate of 13.5 Hz. In the case of 1,000 cSt droplets, both the continuous and dispersed phases were pressurized at 400 mbar, resulting in droplets with an average diameter of 15.5 μm at a rate of 27 Hz. Droplets were collected inside a 15 mL Falcon tube.

### Cell culture

The HL60/S4 cell subline (ATCC Cat# CRL-3306, RRID:CV-CL_II77) was cultured in RPMI 1640 medium with 2 mM L-Glutamine (Thermo Fisher #A1049101) with 1% penicillin and streptomycin (Gibco) and 10% heat-inactivated fetal bovine serum (Sigma Aldrich F4135, lot no. 13C519). Cells were grown at 37 °C, with 5% CO_2_, at densities between 10^5^ – 10^6^ cells per mL with subculturing every 48–72 hours. Cells used for experiments originated all from the same frozen batch after thawing and were measured in 14-26 passages after initial seeding.

### Experimental protocol

Measurements were performed using an AcCellerator device with temperature control (Zellmechanik Dresden GmbH, Germany). This device offers high-speed imaging with online contour analysis through the software ShapeIn 2 (Zellmechanik Dresden). All measurements were performed at a chamber temperature of 25 ± 0.5 °C.

For a measurement, the microfluidic chip was placed on the stage of an inverted microscope (Axio Observer Z1, Zeiss, Germany) and flow was introduced using syringe pumps (neMESyS 290N, Cetoni GmbH, Germany) connected to the chip with FEP tubing (0.0625” OD, 0.03” ID; no. 1520XL, Postnova Analytics). The sheath to sample flow rate ratio was 1.5:1 for oil droplet experiments and 3:1 for all other measurements.

Imaging was done with a pulsed, high power, blue LED (Zellmechanik Dresden) that was synchronized to a CMOS camera (EoSens CL, MC1362, Mikrotron GmbH, Germany) and imaged through a 20× objective (Plan-Apochromat, ×20/0.8; no. 420650-9902, Zeiss). To measure the stresses acting in the channel based on PAAm bead deformation, we used a 40× objective (EC-Plan-Neofluar, ×40/0.75; no. 420360-9900, Zeiss). Tubings for sheath and sample were filled with the respective carrier solution before connecting to the chip and then chips were filled through the sheath inlet using the syringe pump until liquid was pushed out of the sample inlet. Sample suspensions were loaded into the tubing by placing the tubing inside the suspension and setting a negative flow rate of -2 μL/s for the sample. The filled chip was then connected to the filled sample tubing to avoid the introduction of air bubbles into the chip. The samples were pushed into the chip at a flow rate of 0.1 μL/s before setting the measurement flow rate.

#### Oil droplets

accumulated at the top of the emulsion inside the storage container. 1 μL of this concentrated droplet emulsion was pipetted into 50 μL of the glycerol-water mixture and mixed by up-down pipetting before loading into the sample tubing. The silicone oil droplet samples were measured at a flow rate of 0.04 μL/s.

#### PAAm beads

accumulated at the bottom of the storage Eppendorf tube. 1-2 μL of this concentrated bead suspension was pipetted into 98-99 μL of 0.6% MC-PBS and mixed by up-down pipetting before loading into the sample tubing.

#### HL60 cells

were counted on the measurement day and then a volume of the cell culture medium was centrifuged at 200×g for 2 minutes to reach a final concentration of 1-2×10^6^ cells/mL in the cell carrier solution. After centrifugation, the supernatant was taken off and the cell pellet was re-suspended in 100 μL of 0.6% MC-PBS, containing the respective amount of LatB or Nocodazole, and mixed by up-down pipetting. The samples were then stored for an incubation time of 30 minutes at 25 °C inside the heating chamber of the AcCellerator before loading into the sample tubing.

### Data acquisition

Data was recorded using the program ShapeIn 2 (Zellmechanik Dresden). The program analyses the images in real-time by finding a contour based on background subtraction and thresholding.^16^ A number of contour and brightness features are extracted from the image and saved to an hdf5 file.

For a viscoelasticity experiment in the hyperbolic region, objects were recorded in a 680×54 μm region of interest (ROI, see Fig. 1A). The border of the ROI was placed 50 μm after the end of the hyperbolic region (*x* = *L*_*c*_ + 50 μm). The frame rate was adjusted according to the flow rate to capture about 50 events per object flowing through the ROI.

PAAm beads and silicone oil droplets showed optical distortions dependent on the position in the ROI. To correct for these effects, we measured each sample 500 μm before the hyperbolic region with an ROI of 680×102 μm at a flow rate of 0.01 μL/s (see inlet ROI in Fig. 1A). In this region, no extensional field is present and at this flow rate, the influence of shear stresses is negligible. The measured deviations in the deformation can be attributed to optical distortions in the setup. The correction curves are shown in figures S3-S5.

#### Stress measurements

For the stress measurements on PAAm beads inside the hyperbolic region, we used an ROI of 230x30 μm, and for measurements in the inlet or channel 102x34 μm (see Fig. 2A). We used a lager ROI for measurements in the hyperbolic region to measure multiple events per bead, to determine bead velocities and derive extension rates. ROI hyper in figure 2A shows the window used for the strain analysis (see Results section).

**Figure 2:**
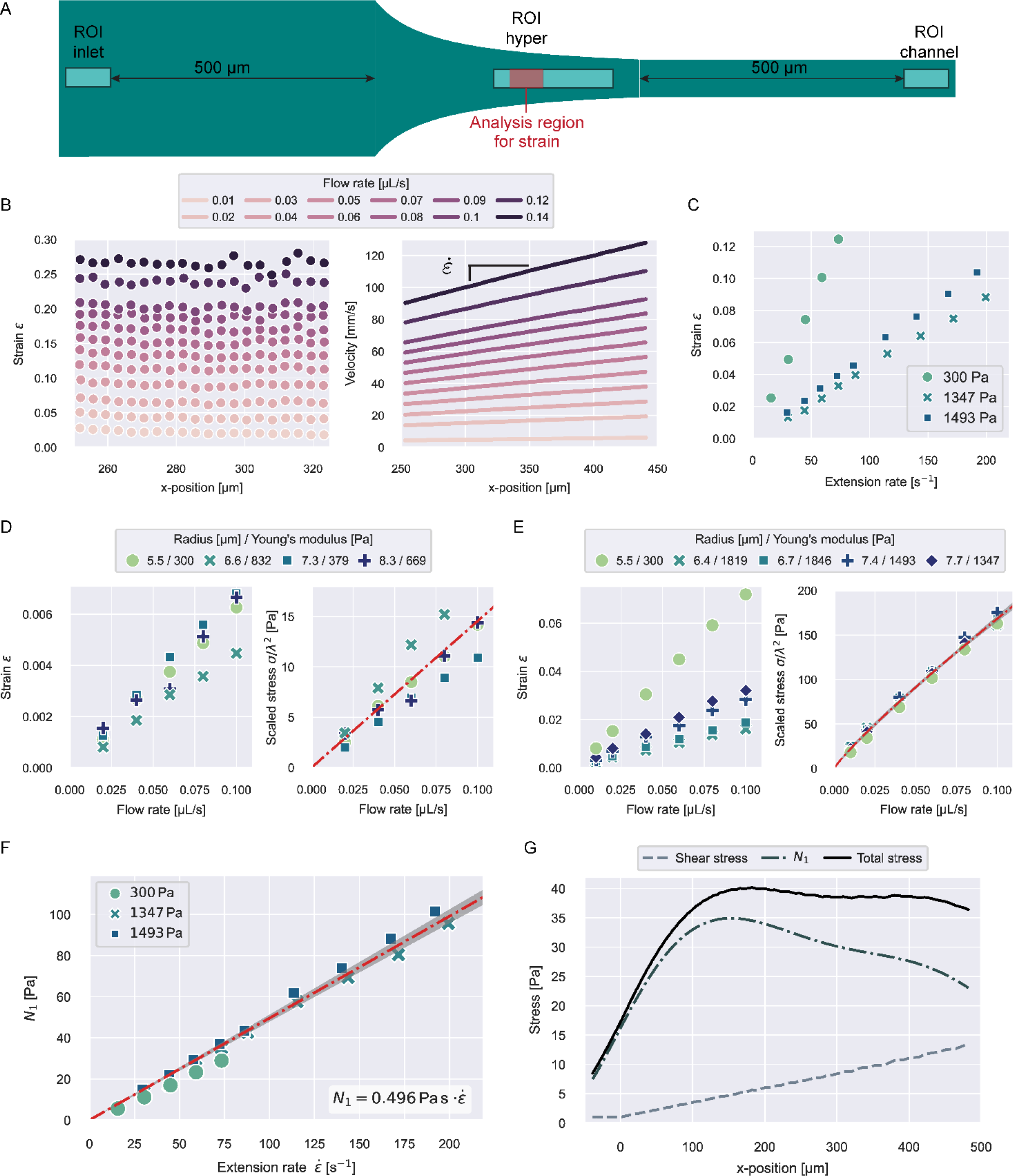
Stress characterization of 0.6% MC-PBS in the hyperbolic region. **A**) ROIs for the different stress measurements. The red region in ROI hyper represents the analysis window for the strain data shown in B. **B**) Strain and velocities of 300 Pa PAAm beads in ROI hyper at different flow rates. **C**) Median strain of PAAm beads with varying stiffness dependent of extension rate in the strain analysis region. **D**) Median strain and apparent shear stress scaled by bead size for different beads in ROI inlet. The line in the scaled stress plot shows a linear fit to the data with 2-σ error band. **E**) Median strain and apparent shear stress scaled by bead size for different beads in ROI channel. The line in the scaled stress plot shows a power law fit to the data with 2-σ error band. **F**) First normal stress difference 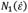 for different beads measured in ROI hyper. The line shows a linear fit to the data with 2-σ error band. **G**) Shear stress, first normal stress difference, and total stress for 830 Pa PAAm beads flowing through the hyperbolic region (see Fig. 1A) at a flow rate of 0.04 μL/s.

### Data analysis

The full analysis pipeline is documented at https://gitlab.gwdg.de/cell-viscoelasticity-in-hyperbolic-channels.

#### Object Tracking

The data from the hdf5 files was tracked to get trajectories for every single object using a custom-made python package called *dctrack*. Velocities and extension rates were computed from the tracked object trajectories. Event times were calculated from the velocity curve ***v***(***x***) in reference to a point ***x***_**0**_ with:

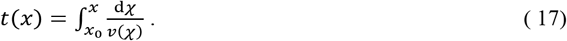

#### Strain computation

Object contours only extend a few pixels in *x* and *y* and the computation of the strain from the raw contour bounding box size is very susceptible to noise in the contour detection. To have a more robust measure, the strain was calculated from the second moments of area of the raw contour over *x, y* defined as:^43^

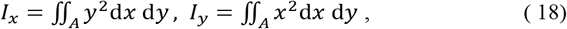

with contour area *A*. The ratio of 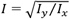 is readily available as a feature in the measurement hdf5 files and known as the *inertia ratio*. For a perfect ellipse or rectangle, *I* is equal to the aspect ratio of the contour. The strain definition in equation (6) can then be rewritten using the inertia ratio:

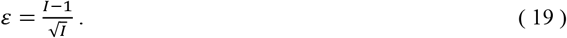

### Data binning

Data curves as function of *x* or time as shown in figures 1-5 were made by creating equal sized bins in *x* and computing the median from all datapoints inside the respective *x* -bin.

**Figure 3:**
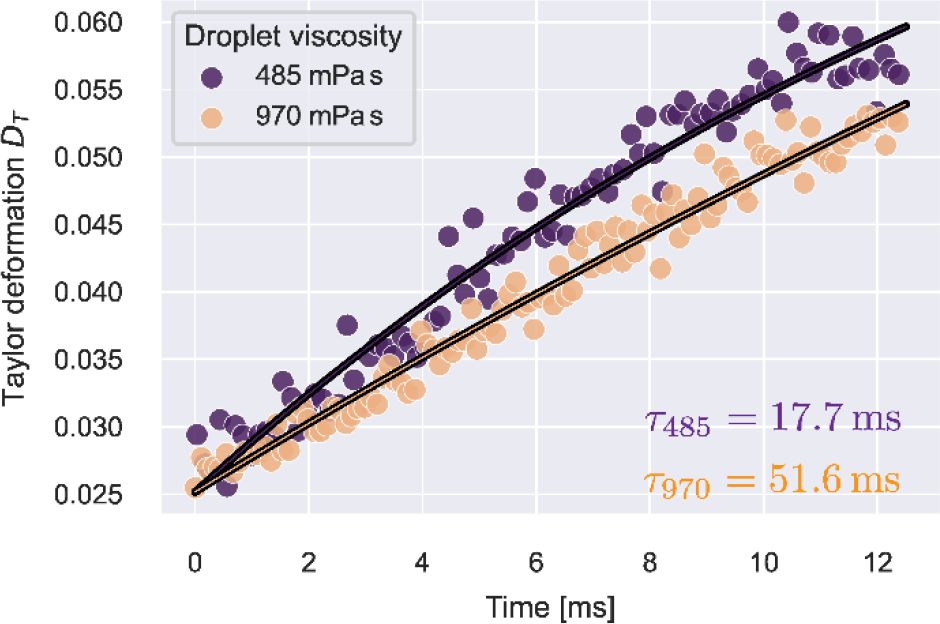
Taylor deformation over time for silicone oil droplets at consistent extension rate. The solid lines illustrate fit curves of equation 27 to the datapoints.

**Figure 4:**
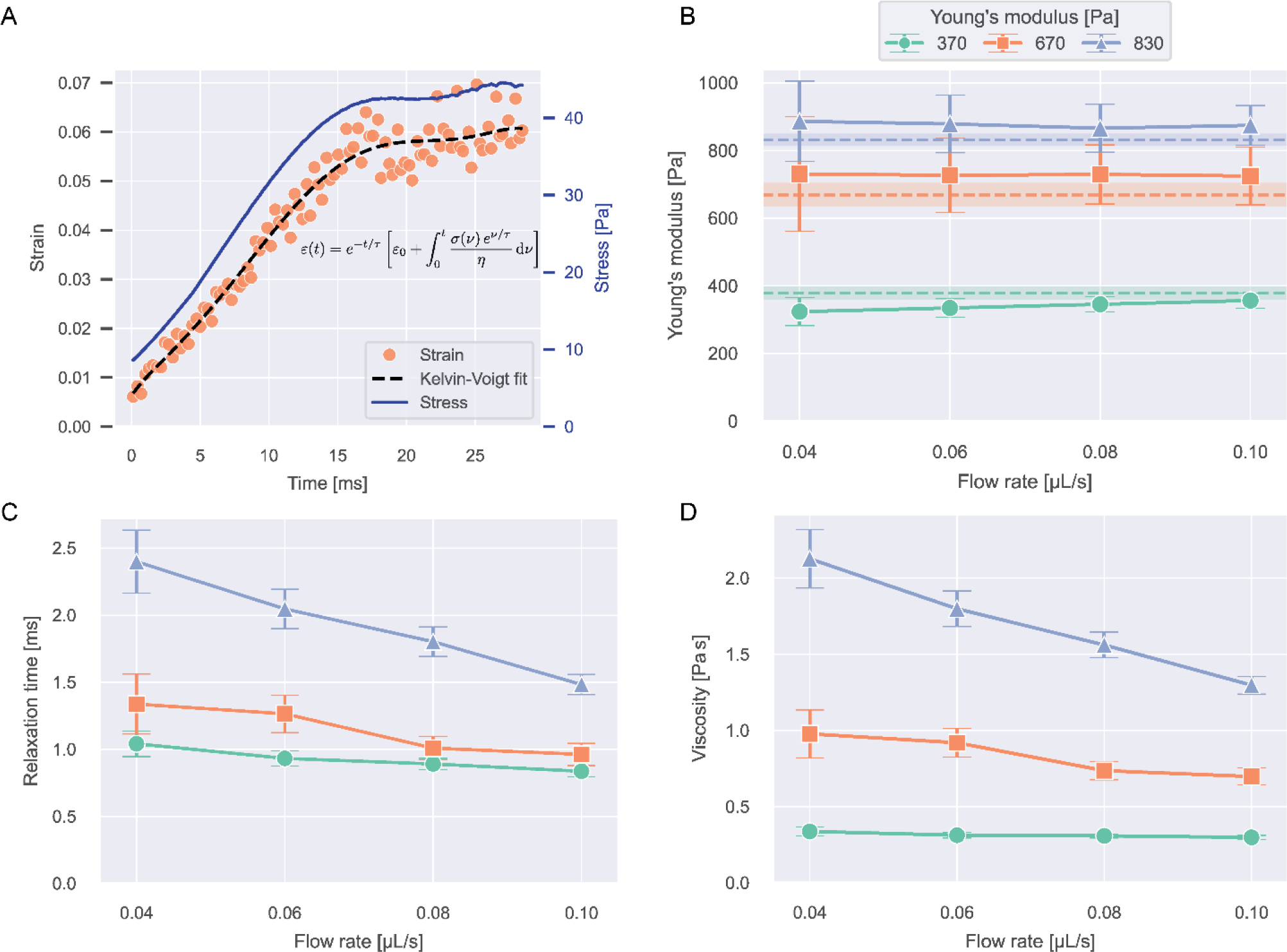
Viscoelasticity of PAAm microgel beads. **A**) Median strains over time and stress curve for 690 Pa beads at 0.04 μL/s flow rate. The dashed line shows the Kelvin-Voigt fit to the datapoints (equation 13). **B**) Young’s moduli resulting from the Kelvin-Voigt fit for different flow rates and bead types. Values show fit result ± standard error of the fit. The dashed lines and shaded regions correspond to the respective mean ± SEM Young’s modulus measured with RT-DC (see Fig. S1). **C**) Relaxation times and **D**) Viscosities of the beads resulting from the fit.

**Figure 5:**
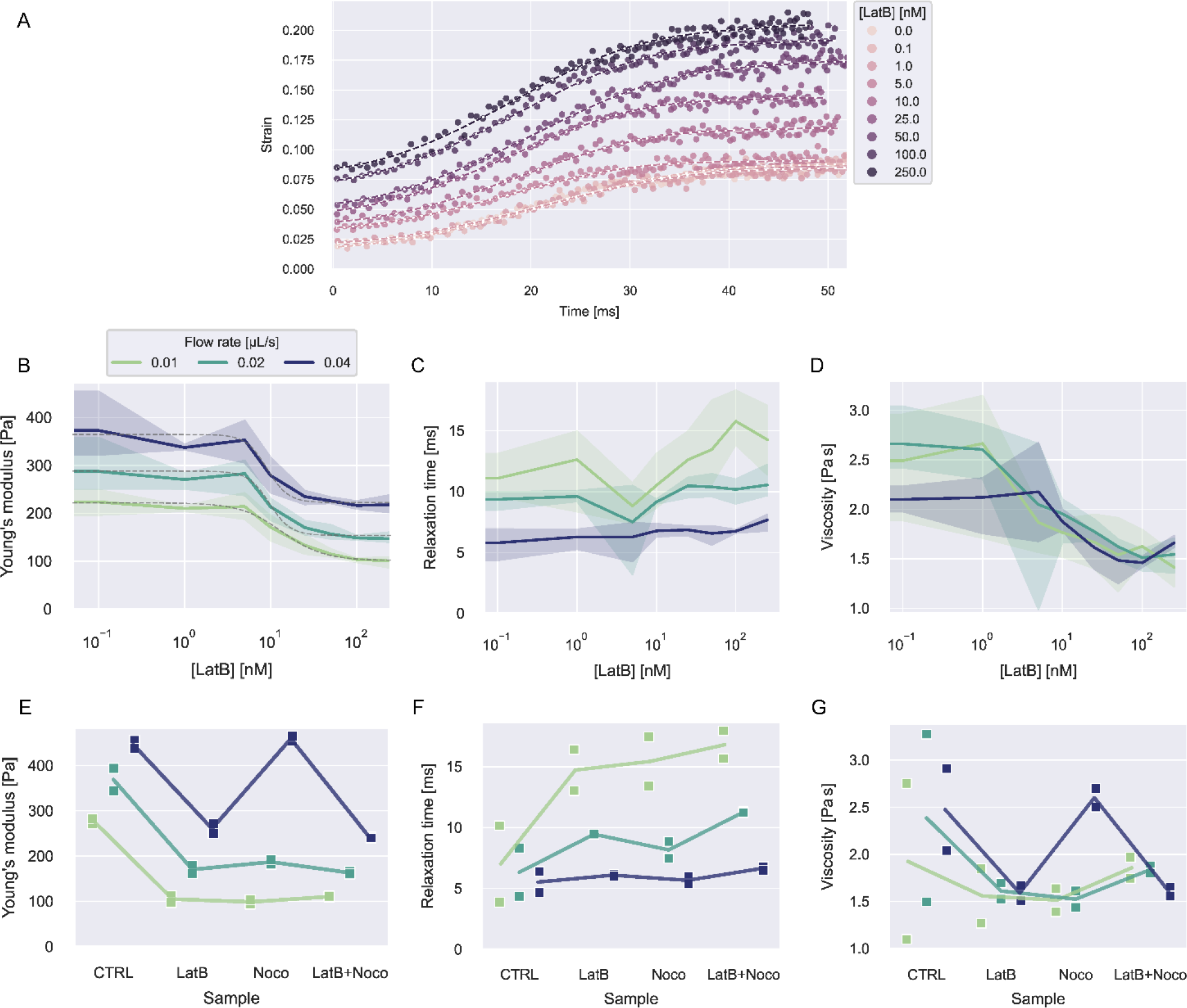
Viscoelasticity of HL60 cells treated with LatB and Nocodazole. **A**) Exemplary strain curves over time and fit curves at different LatB concentration for one experiment date at flow rate 0.02 μL/s. **B-D**) Young’s moduli, relaxation times, and viscosities dependent on LatB concentration at different flow rates. The solid lines represent the mean value at each concentration and the error bands represent the full range of datapoints. The dashed gray lines in B show fits of a log-logistic growth function to the data (see SI text). **E-G**) Young’s moduli, relaxation times, and viscosities for LatB, Nocodazole or combined LatB+Nocodazole treatment. The solid lines represent the mean of the measurements at each condition.

## Results and Discussion

### Characterizing stresses in the hyperbolic region

The first normal stress differences of 0.6 w% MC-PBS in extensional flow were not reported before. We measured the stresses acting in the hyperbolic region using PAAm microgel beads. The Young’s modulus, *E*, of the beads was measured using RT-DC (see SI text and Fig. S1). The first normal stress difference was then deducted from the strain measured in the hyperbolic region:

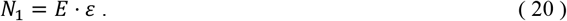

To find a relation of *N*_1_ and the extension rate 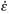, beads were measured at different flow rates and the ROI was placed 225 μm after the start of the hyperbolic region (see “ROI hyper” in Fig. 2A). This region was chosen to ensure the beads were measured in a zone with a consistent extension rate and to give the beads enough time to comply to the stress. Figure 2B shows the median strain and velocity curves of beads with *E* = 300 Pa. The analysis region was restricted to 250 < *x* < 325 μm. This was done to minimize the influence of shear stresses on the bead deformation when getting closer to *x* = *L*_*c*_. It can be seen that the strain is constant over *x* in the chosen analysis region for all flow rates.

The velocity curves were analyzed from the full ROI width of 230 μm. Velocities were computed for single beads from the displacement change between frames. The velocities were then assigned to the second frame and no velocity data is recorded for the start of the ROI. In figure 2B we show the velocities for 250 μm < *x*. The extension rates were computed from the slope of a linear fit to the velocity data.

The resulting median strains and extension rates for three bead types of different stiffness are shown in figure 2C. The further analysis was restricted to small strains with *ε* < 0.125 to stay within the linear elastic regime. The strain values were corrected for influences of optical distortions from the instrument (see Fig. S2).

Typical object diameters for our experiments were in the range of 10-20 μm. At a channel height of 30 μm, the influence of shear stresses on the deformation cannot be neglected. To quantify this influence, we measured the deformation of the beads 500 μm before entering and after leaving the hyperbolic region (see “ROI inlet” and “ROI channel” in Fig. 2A). The resulting strains are shown figure 2D,E. The apparent shear stress *σ*_*s*_ can then be calculated similarly to equation (20).

The shear stress on an object inside the channel is size-dependent because shear stresses increase closer to the channel walls. We found that scaling the apparent stresses quadratically by size resulted in the datapoints collapsing on one line (see Fig. 2D,E). We performed the scaling with the confinement λ defined by:

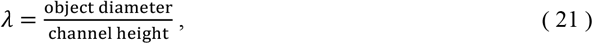

The scaled stress *σ*_λ_ is defined by:

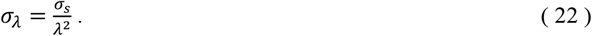

The data for *σ*_λ_(*Q*) at the inlet was best described by a linear function while it showed shear-thinning power law behavior inside the channel (Fig. 2D,E). Fits to the data resulted in the following functions for the shear influence:

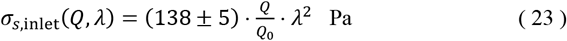

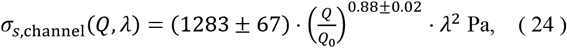

with *Q*_0_ = 1 μL/s.

Since the velocity increases approximately linearly in the hyperbolic region, we can assume the apparent shear stresses will also increase linearly. The apparent shear at any point *xx* in the contraction can then be calculated with:

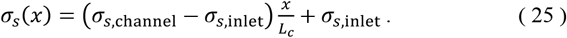

Considering the influence of the shear stress, the first normal stress difference can then be computed from the data in figure 2C, equation 20 and correcting for *σ*_*s*_:

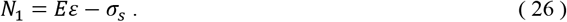

The resulting values for *N*_1_ as function of the extension rate are shown in figure 2F. It can be seen that the datapoints from different bead types all collapse on one line. A linear function was fitted to the data which resulted in an intercept close to zero. We fixed the intercept at zero because the solution should be stress-free for 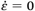. A linear relation of *N*_1_ and 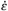 corresponds to a constant extensional viscosity for the 0.6 w% MC-PBS solution up to 200 *ss*^−1^ with *η*_*E*_ = (496 ± 8) mPa s. A similar behavior for aqueous MC solutions with addition of salt was described by Micklavzina et al.^25^ It is noticeable that the extensional viscosity of the solution is much higher than the zero shear viscosity, which is reported at 30 mPa s, highlighting the need for distinct measurements in extension to accurately determine the stresses by MC-solutions.

Finally, the total stress on an object at any position *x* inside the hyperbolic region can be calculated from the extension rate 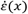, the flow rate, and the initial object diameter:

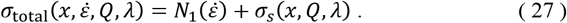

The stress curves for beads with an apparent Young’s modulus of 832 Pa and a radius of 6.6 μm at a flow rate of 0.04 μL/s are shown in figure 2G. It can be seen that the total stress will stay approximately constant after about 150 μm.

### Mechanics of silicone oil droplets

To verify that our approach is sensitive to detect different relaxation times, we measured silicone oil droplets with different viscosities. The lower viscosity droplets had a dynamic viscosity of 485 mPa s and the higher viscosity sample 970 mPa s, according to manufacturer.

Droplets were analyzed by the Taylor deformation as introduced in equation 7. The full strain and extension rate curves as function of *x* are shown in figure S3. A consistent extension rate was achieved after *x*_0_ = 110 μm. The droplet deformation was analyzed in the range 110 μm < *x* < 390 μm. The upper limit for *x* was chosen to avoid strong effects of the shear stress influencing the deformation. The resulting Taylor strain over time, where *t*(*x*_0_) = 0, is show in figure 3.

According to equation 10, the data follows an exponential and will equilibrate at *D*_*∞*_. At *t* = 0, the data shows a pre-deformation *D*_0_ caused by stresses acting on the droplets before reaching *x*_0_. With these, the deformation curve will have the form:

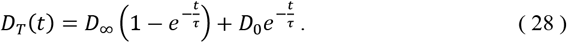

Equation 28 was fitted to the data to get *D*_*∞*_ and the relaxation time, τ. We found τ(485 mPa s) = 18 ± 4 ms and τ(970 mPa s) = 52 ± 25 ms, highlighting that we are sensitive to different time scales of droplet deformation. Error margins show standard errors of the fit. τ(970 mPa s) has a large error margin of almost 50%. This large error can be explained because the estimated relaxation time is much larger than the observation time of about 12 ms, which leads to inaccuracies when fitting the exponential.

Utilizing the analysis in equations 8 – 11, we can determine the interfacial tensions and viscosities of the droplet systems. For this, the extension rate was taken as a constant average, calculated from all datapoints in the analysis region. For the 485 mPa s droplets, we get an interfacial tension of *γ* = 1.0 ± 0.1 mN/m and the viscosity *η*_drop_ = 552 ± 73 mPa s, which results in the expected viscosity of the silicone oil, within error margins. For 970 mPa s droplets we get *γ* = 0.6 ± 0.2 mN/m and *η*_drop_ = 711 ± 212 mPa s. The deviation of the resulting viscosity from the expected value can be explained by the large error margin of the relaxation time. Both emulsified droplet systems show a small interfacial tension of around 1 mN/m. Similar values for such interfaces have been reported before.^44^

### Viscoelasticity of PAAm-microgel beads

The strain curves of PAAm-microgel beads with Young’s moduli 370, 690, and 830 Pa were measured at flow rates from 0.04-0.10 μL/s and analyzed for their viscoelastic behavior with the Kelvin-Voigt model explained in equation 13. Object times were calculated in reference to *x*_0_ = −40 μm. The analysis was restricted to values for −40 μm < *x* < 490 μm. Strain curves were corrected for optical distortions as shown in figure S4. An example stress and strain curve as function of time with Kelvin Voigt fit for 690 Pa beads at 0.04 μL/s is shown in figure 4A. The data for all conditions are shown in figure S4.

Figure 4B-D shows the Young’s moduli, relaxation times and viscosities resulting from the fit. The error bars indicate standard errors of the fit.

Young’s moduli stayed constant over different flow rates for all bead types and within error margins, the expected values for all conditions could be recaptured, showing that our approach is able to accurately measure object stiffness. Relaxation times were longer for stiffer beads and generally showed a decrease with increasing flow rate. The same behavior was observed for the bulk viscosities of the beads.

With relaxation times on the scale of millisecond, the beads deform almost instantaneously, which can be seen when comparing the strain and stress curves. This implies near-elastic behavior, which is expected for microgels at these timescales.^10,29^

### Viscoelasticity of HL60 cells after LatB and Nocodazole treatment

To investigate how actin is contributing to the viscoelastic properties of HL60 cells, we treated the cells with Latrunculin B (LatB) at concentrations between 0 – 250 nM dissolved in the 0.6% MC-PBS solution. LatB is known to induce cell softening by inhibiting the polymerization of F-actin.^15,45,46^

Object times were calculated in reference to *x*_0_ = −50 μm. The analysis was restricted to values for −50 μm < *x* < 480 μm. Cells were measured at flow rates 0.01, 0.02, and 0.04 μL/s and biological triplicates from different culturing passages were measured for every concentration.

In contrast to beads and oil droplets, the cells showed no position-dependent deformation in the inlet but constant strain offsets, that could not be explained solely by the influence of the shear stress (see Fig. S5). To account for this offset, we modified equation 13 by addition of a constant strain offset *ε*_offset_:

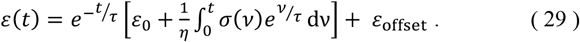

This function was fitted to the strain function in time. The resulting curves from one experiment date at 0.02 μL/s are shown in figure 5A. All strain and corresponding stress curves are shown in figure S6.

Figure 5B-C shows the resulting viscoelastic parameters. The Young’s moduli at all flow rates show a decrease with increasing LatB concentration that follows a log-logistic growth function (see SI text and table S2). This behavior has been described before for Latrunculin-treated cells.^3,15^ Young’s moduli increased with increasing flow rate for all concentrations. The apparent increase of elastic moduli of other cell types when measured at faster time scales is widely described in the literature.^3,21,23^

Relaxation times showed a slight increase with LatB concentration. Measuring at higher flow rates resulted in faster relaxation times. The relaxation times vary in a range from about 5-15 ms. Similar values have been reported using other microfluidic techniques.^5,23^

The resulting bulk viscosities show similar behavior to the Young’s modulus, because the relaxation times only show small changes in comparison. In contrast to the other two parameters, the viscosities show no clear trend with increasing flow rate.

To investigate how microtubuli contribute to the viscoelastic properties, we treated the cells with Nocodazole to disrupt the microtubuli polymerization. We used a concentration of 1 μM Nocodazole dissolved in 0.6% MC-PBS. Additionally, we treated cells at 250 nM LatB and a combination of both treatments. We performed measurement on biological duplicates from different passages for these treatments.

The resulting strain, stress, and fit curves are shown in figure S7 and the viscoelastic parameters are shown figure 5E-G. We would like to point out that the LatB data in these results comes from different experiments than the data shown in figure 5B-C. It can be seen that Nocodazole in addition to LatB had nearly no further influence on the cells’ Young’s modulus. Treatment with Nocodazole alone had a similar effect as LatB when measured at 0.01 and 0.02 μL/s but at a flow rate of 0.04 μL/s, the cells even showed a slight increase in the Young’s modulus.

The treatment effect on the relaxation times was flow rate-dependent. At 0.01 μL/s, the three treatments showed a similar increase in the relaxation time compared to the control condition. At 0.02 μL/s, this effect was less pronounced and at 0.04 μL/s, the treatment had nearly no influence on the relaxation times.

The viscosities showed no clear trend with the treatments. For 0.01 and 0.02 μL/s, the treatments lead to a reduction of the mean viscosity but the spread in the data is too large to make definite statements. At 0.04 μL/s, the viscosity follows the behavior of the Young’s modulus, as the relaxation time was hardly affected by the treatments.

### Discussion

Our platform was well-suited for accurately measuring the stresses applied by 0.6% MC-PBS in extensional flow. We observed a constant extensional viscosity of 0.5 Pa s up to an extension rate of 200 s^-1^. This aligns with findings from previous studies on MC solutions with added salt.^25^ Notably, the extensional viscosities measured were more than 10 times larger than shear viscosities at comparable shear rates. This emphasizes the necessity to independently assess the stresses exerted by polymer solutions in extensional flow, separate from shear stresses.^25,26^

Measurements on silicone oil droplets showcased the platform’s sensitivity to changes in relaxation times, allowing for the determination of interfacial tension and droplet viscosities. Challenges were encountered in accurately determining viscosity for droplets with high viscosity due to observation time limitations. This emphasizes the importance of carefully choosing flow rates to guarantee enough time for deformation.

In experiments on PAAm beads, our platform effectively recaptured expected Young’s moduli. The rapid adaptation of strain to stress curves in beads suggested near-elastic behavior. We observed that stiffer beads had longer relaxation times. This behavior was rather unexpected for deformable particles, where higher stiffness often leads to faster relaxation.^21^ A possible explanation for this effect could be the poroelastic nature of the PAAm microgel.^10,29,47^ Stiffer beads are made of a more densely connected polymer network, leading to smaller pores. Smaller pores cause a smaller permeability of the material for fluid flow. This increased resistance for fluids to move through the microgels could lead to longer relaxation times.^10^

A viscoelastic response of the microgel can also be influenced by rearrangements of polymers chains or cross-links in the network. This would normally not be expected for PAAm hydrogels but defects in the bead production can lead to such rearrangements when stress is applied.^29^

We observed a decrease of bead relaxation times and viscosities with increasing flow rate. A similar effect was reported by Abuhattum et al., when measuring PAAm hydrogel viscosity with increasing AFM indentation velocity.^10^ As the timescale of deformation in the poroelastic hydrogel primarily depends on the rate at which liquid moves through the expanded elastic polymer network, the observed effect may be attributed to an accelerated fluid flow under increased stresses on the material. This effect may be amplified by the shear thinning of the MC-PBS solution.^24^

Examining HL60 cell behavior under LatB treatment revealed a decrease in Young’s modulus with increasing LatB concentration, consistent with prior studies.^3,15,45,46^ Intriguingly, at higher flow rates, cells exhibited increased stiffness irrespective of LatB concentration, suggesting an actin-independent effect. Similar stiffening effects were observed with additional nocodazole treatment.

Higher elastic moduli in measurements at shorter time scales or under greater stresses have been previously reported.^3,21,48^ Another possible explanation for our observation is that cells do not behave as linear elastic materials but rather follow a hyperelastic model.^19,49^ This can lead to an overestimation of the stiffness at larger deformations.

Nocodazole treatment led to a decrease in Young’s modulus at slower flow rates and a slight increase at the highest flow rate. The literature is not clear on the effect of nocodazole on cell stiffness: Golfier et al. reported on reduced cell deformability with nocodazole treatment, while Liu et al. reported a decreasing Young’s modulus.^46,50^ Besides inducing microtubuli disassembly, nocodazole is known to increase cell contractility by releasing GEF-H1 from microtubuli, which consequently leads to RhoA and Rock activation and the formation of actin stress fibers.^51^ This can lead to stiffening of cells besides the depletion of microtubuli. Our results suggest that this interplay of microtubule disassembly and increased cell contractility might have time scale dependent effects on cell stiffness. The effect of LatB and/or nocodazole treatment on relaxation times was most pronounced at slower flow rates. Cells are known to exhibit more fluid-like behavior when deformed over a longer time scale.^4,21,48,52^ At slower flow rates, the cells have more time to adapt to the stress in the channel. The viscous properties only become apparent when cells get stretched over a longer period. This highlights that flow rates need to be chosen carefully to reveal the viscous properties of cells.

## Conclusions

Our microfluidic platform, tailored for high-throughput viscoelastic characterization at up to 100 cells per second, has provided profound insights into the behaviors of complex fluid systems and soft deformable particles.

Experiments on droplets, PAAm microgel beads and HL60 cells showed that our platform is sensitive to changes in material stiffness and relaxation times. To investigate the influence of filamentous actin and microtubuli on cell viscoelastic properties, we treated HL60 cells with Latranculin B, Nocodazole, or both. Our results highlight that the measurement time scales affect the viscoelastic response to the treatments. Our current analysis requires a cautious estimation of shear and tensile stresses acting in the microchannel. The influence of shear stresses, mainly resulting from the velocity profile between bottom and top wall in the channel, could be minimized by increasing the height of the channels. This would require an additional focusing step to focus the cells in the center of the channel height.^53^

Furthermore, the channel profile could be improved to have a better-defined development of the extension rate and a true constant extension rate in the hyperbolic region.^54^ This would help to get simpler stress estimates in the hyperbolic region and would lead to more straight-forward approaches for analyzing the stress-strain relationship.

Our results illustrate the great potential of hyperbolic microchannel systems for high-throughput measurements of single cell viscoelasticity with a wide range of applications in cell research or diagnostics.

## Data availability statement

Data and analysis scripts to recreate the results presented here can be found at gitlab: https://gitlab.gwdg.de/cell-viscoelasticity-in-hyperbolic-channels. (Persistent identifier (PID): 21.11101/0000-0007-FC23-6; see https://www.pidconsortium.net)

## Author Contributions

Conceptualization: FR, JG; Data curation: FR; Formal analysis: FR; Investigation: FR, Methodology: FR, RG, SG; Project administration: JG; Resources: RG, SG, JG; Software: FR; Supervision: SG, JG; Validation: FR; Visualization: FR; Funding acquisition: SG, JG; Writing – original draft: FR; Writing – editing and review: all authors

## Acknowledgements

We want to thank Benedikt Hartmann, Shada Abuhattum, and Sebastian Bohle for helpful discussion regarding this study. Additional thanks to Benedikt Hartmann and Eoghan O’Connel for help with *dctrack*. Special regards to Cornelia Liebers, Manuela Hauke and Christine Schweitzer for taking care of cell culture. Further thanks to Cornelia Liebers for help with production of the MC-PBS solutions. We want to thank Parth Patel, who produced the microfluidic chips for this work. We would like to acknowledge the assistance of ChatGPT3.5, a language model developed by OpenAI, in refining the language of this article. The authors acknowledge the financial support through the base funding of the Max Planck Society to JG. Open Access funding provided by the Max Planck Society. RG and SG receive funding from the European Unions Horizon 2020 research and innovation programs No. 953121 (project FLAMIN-GO).

## Notes and references

1. Zak, A. et al. Rapid viscoelastic changes are a hallmark of early leukocyte activation. Biophys J, 2021, 120, 1692–1704.

2. Abidine, Y., Giannetti, A., Revilloud, J., Laurent, V. M. & Verdier, C. Viscoelastic properties in cancer: From cells to spheroids. Cells, 2021, 10, 1704.

3. Gerum, R. et al. Viscoelastic properties of suspended cells measured with shear flow deformation cytometry. Elife, 2022, 11.

4. Dupire, J., Puech, p. H., Helfer, E. & Viallat, A. Mechanical adaptation of monocytes in model lung capillary networks. Proc Natl Acad Sci U S A, 2020, 117, 14798–14804.

5. Fregin, B. et al. High-throughput single-cell rheology in complex samples by dynamic real-time deformability cytometry. Nat Commun, 2019, 10, 415.

6. Abuhattum, S. et al. Adipose cells and tissues soften with lipid accumulation while in diabetes adipose tissue stiffens. Sci Rep, 2022, 12, 10325.

7. Hochmuth, R. M. Micropipette aspiration of living cells. J Biomech, 2000, 33, 15–22.

8. Efremov, Y. M., Okajima, T. & Raman, A. Measuring viscoelasticity of soft biological samples using atomic force microscopy. Soft Matter, 2020, 16, 64–81.

9. Staunton, J. R., Doss, B. L., Lindsay, S. & Ros, R. Correlating confocal microscopy and atomic force indentation reveals metastatic cancer cells stiffen during invasion into collagen I matrices. Sci Rep, 2016, 6, 19686.

10. Abuhattum, S. et al. An explicit model to extract viscoelastic properties of cells from AFM force-indentation curves. iScience, 2022, 25, 104016.

11. Guck, J. et al. The Optical Stretcher: A Novel Laser Tool to Micromanipulate Cells. Biophys J, 2001, 81, 767–784.

12. Guck, J. et al. Optical Deformability as an Inherent Cell Marker for Testing Malignant Transformation and Metastatic Competence. Biophys J, 2005, 88, 3689–3698.

13. Fuhs, T. et al. Rigid tumours contain soft cancer cells. Nat Phys, 2022, 18, 1510–1519.

14. Huster, C. et al. Stretching and heating cells with light—nonlinear photothermal cell rheology. New J Phys, 2020, 22, 085003.

15. Urbanska, M. et al. A comparison of microfluidic methods for high-throughput cell deformability measurements. Nat Methods, 2020, 17, 587–593.

16. Otto, O. et al. Real-time deformability cytometry: on-the-fly cell mechanical phenotyping. Nat Methods, 2015, 12, 199–202.

17. Gossett, D. R. et al. Hydrodynamic stretching of single cells for large population mechanical phenotyping. Proc Natl Acad Sci USA, 2012, 109, 7630–7635.

18. Wittwer, L. D., Reichel, F. & Aland, S. Numerical simulation of deformability cytometry: Transport of a biological cell through a microfluidic channel. in Modeling of Mass Transport Processes in Biological Media 33–56 (Elsevier, 2022). doi:10.1016/B978-0-323-85740-6.00010-8.

19. Wittwer, L. D., Reichel, F., Müller, P., Guck, J. & Aland, S. A new hyperelastic lookup table for RT-DC. Soft Matter, 2023, 19, 2064–2073.

20. Abuhattum, S., Kuan, H.-S., Müller, P., Guck, J. & Zaburdaev, V. Unbiased retrieval of frequency-dependent mechanical properties from noisy time-dependent signals. Biophys Rep, 2022, 2, 100054.

21. Kollmannsberger, P. & Fabry, B. Linear and Nonlinear Rheology of Living Cells. Annu Rev Mater Res, 2011, 41, 75–97.

22. Piergiovanni, M. et al. Deformation of leukaemia cell lines in hyperbolic microchannels: investigating the role of shear and extensional components. Lab Chip, 2020, 20, 2539–2548.

23. Armistead, F. J., Gala De Pablo, J., Gadêlha, H., Peyman, S. A. & Evans, S. D. Cells Under Stress: An Inertial-Shear Microfluidic Determination of Cell Behavior. Biophys J, 2019, 116, 1127–1135.

24. Büyükurganci, B. et al. Shear rheology of methyl cellulose based solutions for cell mechanical measurements at high shear rates. Soft Matter, 2023, 19, 1739–1748.

25. Micklavzina, B. L., Metaxas, A. E. & Dutcher, C. S. Microfluidic rheology of methylcellulose solutions in hyperbolic contractions and the effect of salt in shear and extensional flows. Soft Matter, 2020, 16, 5273–5281.

26. Kim, S. G., Ok, C. M. & Lee, H. S. Steady-state extensional viscosity of a linear polymer solution using a differential pressure extensional rheometer on a chip. J Rheol (N Y N Y), 2018, 62, 1261–1270.

27. Lee, S. S., Yim, Y., Ahn, K. H. & Lee, S. J. Extensional flow-based assessment of red blood cell deformability using hyperbolic converging microchannel. Biomed Microdevices, 2009, 11, 1021–1027.

28. Faustino, V. et al. A Microfluidic Deformability Assessment of Pathological Red Blood Cells Flowing in a Hyperbolic Converging Microchannel. Micromachines, 2019, 10, 645.

29. Girardo, S. et al. Standardized microgel beads as elastic cell mechanical probes. J Mater Chem B, 2018, 6, 6245–6261.

30. Ober, T. J., Haward, S. J., Pipe, C. J., Soulages, J. & McKinley, G. H. Microfluidic extensional rheometry using a hyperbolic contraction geometry. Rheol Acta, 2013, 52, 529–546.

31. Macosko, C. W. Rheology: Principles, Measurements, and Applications. (Wiley-VCH Verlag, 1994).

32. Taylor, G. I. The formation of emulsions in definable fields of flow. Proceedings of the Royal Society of London. Series A, 1934, 146, 501–523.

33. Rallison, J. M. The deformation of small viscous drops and bubbles in shear flows. Annu Rev Fluid Mech, 1984, 16, 45–66.

34. Villone, M. M., Nunes, J. K., Li, Y., Stone, H. A. & Maffettone, p. L. Design of a microfluidic device for the measurement of the elastic modulus of deformable particles. Soft Matter, 2019, 15, 880–889.

35. Roscoe, R. On the rheology of a suspension of viscoelastic spheres in a viscous liquid. J Fluid Mech, 1967, 28, 273–293.

36. Cabral, J. T. & Hudson, S. D. Microfluidic approach for rapid multicomponent interfacial tensiometry. Lab Chip, 2006, 6, 427.

37. Choi, Y.-S., Seo, K.-W. & Lee, S.-J. Lateral and cross-lateral focusing of spherical particles in a square microchannel. Lab Chip, 2011, 11, 460–465.

38. Di Carlo, D., Edd, J. F., Humphry, K. J., Stone, H. A. & Toner, M. Particle Segregation and Dynamics in Confined Flows. Phys Rev Lett, 2009, 102, 094503.

39. Bruus, H. Governing Equations in Microfluidics. Microscale Acoustofluidics, 2014, 1–28.

40. Cheng, N.-S. Formula for the Viscosity of a Glycerol-Water Mixture. Ind Eng Chem Res, 2008, 47, 3285–3288.

41. Volk, A. & Kähler, C. J. Density model for aqueous glycerol solutions. Exp Fluids, 2018, 59, 75.

42. Kim, S. C., Sukovich, D. J. & Abate, A. R. Patterning microfluidic device wettability with spatially-controlled plasma oxidation. Lab Chip, 2015, 15, 3163–3169.

43. Herbig, M. et al. Real-Time Deformability Cytometry: Label-Free Functional Characterization of Cells. in Methods in Molecular Biology vol. 1678 347–369 (Humana Press, New York, NY, 2018).

44. Posocco, P. et al. Interfacial tension of oil/water emulsions with mixed non-ionic surfactants: comparison between experiments and molecular simulations. RSC Adv, 2016, 6, 4723–4729.

45. Wakatsuki, T., Schwab, B., Thompson, N. C. & Elson, E. L. Effects of cytochalasin D and latrunculin B on mechanical properties of cells. J Cell Sci, 2001, 114, 1025–1036.

46. Golfier, S. et al. High-throughput cell mechanical phenotyping for label-free titration assays of cytoskeletal modifications. Cytoskeleton, 2017, 74, 283–296.

47. Kalcioglu, Z. I., Mahmoodian, R., Hu, Y., Suo, Z. & Van Vliet, K. J. From macro-to microscale poroelastic characterization of polymeric hydrogels via indentation. Soft Matter, 2012, 8, 3393.

48. Fabry, B. et al. Scaling the Microrheology of Living Cells. Phys Rev Lett, 2001, 87, 148102.

49. Müller, S. J. et al. A hyperelastic model for simulating cells in flow. Biomech Model Mechanobiol, 2021, 20, 509–520.

50. Liu, Y., Mollaeian, K., Shamim, M. H. & Ren, J. Effect of F-actin and Microtubules on Cellular Mechanical Behavior Studied Using Atomic Force Microscope and an Image Recognition-Based Cytoskeleton Quantification Approach. Int J Mol Sci, 2020, 21, 392.

51. Chang, Y.-C., Nalbant, P., Birkenfeld, J., Chang, Z.-F. & Bokoch, G. M. GEF-H1 Couples Nocodazole-induced Microtubule Disassembly to Cell Contractility via RhoA. Mol Biol Cell, 2008, 19, 2147–2153.

52. Trepat, X. et al. Universal physical responses to stretch in the living cell. Nature, 2007, 447, 592–595.

53. Zhao, Y., Li, Q. & Hu, X. Universally applicable three-dimensional hydrodynamic focusing in a single-layer channel for single cell analysis. Anal Methods, 2018, 10, 3489–3497.

54. Zografos, K., Pimenta, F., Alves, M. A. & Oliveira, M. S. N. Microfluidic converging/diverging channels optimised for homogeneous extensional deformation. Biomicrofluidics, 2016, 10.

