## Supplementary Information for "High-throughput viscoelastic characterization of cells in hyperbolic microchannels"

### Supplementary information to article: Viscoelastic characterization of cells at high-throughput in hyperbolic microchannels

#### Interfacial tension and droplet viscosity from hyperbolic channel measurements

In the main text, equations 7-11 introduce a formalism how to calculate interfacial tensions and relaxation times from the deformation $D_T$ of a droplet at a constant extension rate $\dot{\varepsilon}$. This introduces two equations (8 and 11) to compute the interfacial tension from the measurement parameters. By equating equation 8 and 11, one can derive an equation to calculate the droplet viscosity $\eta_{\mathrm{drop}}$ from the extension rate, the viscosity of the carrier solution $\eta_0$, the droplet relaxation time $\tau$, and the steady state deformation $D_\infty$:

$$\frac{5\dot{\varepsilon}\tau}{D_\infty} = \hat{\eta}(2\hat{\eta} + 3)$$

$$\Rightarrow 0 = \hat{\eta}^2 + \frac{3}{2}\hat{\eta} - \frac{5\dot{\varepsilon}\tau}{2D_\infty}$$

$$\Rightarrow \eta_{\mathrm{drop}} = \eta_0 \left(\sqrt{\frac{9}{16} + \frac{5}{2}\frac{\dot{\varepsilon}\tau}{D_\infty}} - \frac{3}{4}\right). \qquad \text{(S1)}$$

$D_\infty$ and $\tau$ result from an exponential fit to the deformation data (equation 28). The extension rate $\dot{\varepsilon}$ is derived from the droplet velocity. The interfacial tension can then be calculated with either equation 8 or 11.

#### Real-time deformability cytometry

The stiffness of PAAm beads used in this study was determined by measurements with real-time deformability cytometry (RT-DC). RT-DC was introduced in 2015 by Otto et al.[1] To determine the Young's modulus, we used the model described by Wittwer et al.[2] and viscosity model for the 0.6% MC-PBS from Büyükurgancı et al.[3]

In brief, we used the same AcCellerator setup (Zellmechanik Dresden) for the RT-DC measurements as for the experiments in hyperbolic channels. The microfluidic chip had a measurement region with cross-section of 30×30 µm, where the deformed beads were recorded. The resulting Young's moduli for each bead type are shown in figure S1 and the median Young's moduli are listed in table S1.

#### Derivation of hyperbolic channel profile

Equation 15 introduces the construction formula for the hyperbolic contraction with the aim of achieving a constant extension rate along the flow centerline. The centerline velocity is calculated according to equation 14. The derivate $\partial u_0/\partial x$ reads:

$$\frac{\partial u_0}{\partial x} = \frac{3Q}{2H}\frac{1}{(w(x) - 0.63H)^2} \cdot \frac{\partial w}{\partial x} \equiv \dot{\varepsilon}$$

This leads to the following differentials:

$$\frac{2H\dot{\varepsilon}}{3Q}\mathrm{d}x = \frac{\mathrm{d}w}{w(x) - 0.63H}. \qquad \text{(S2)}$$

With $A = \frac{2H\dot{\varepsilon}}{3Q}$, integration leads to the following relation:

$$w(x) = 0.63H - \frac{1}{Ax + C_1}, \qquad \text{(S3)}$$

With an integration constant $C_1$. By Fixing $w(0) = w_c$, one gets equation 15 presented in the main text. At the contraction length $L_c$, Equation 15 becomes:

$$w(L_C) = w_u = 0.63H - \left(AL_c - \frac{1}{0.63H - w_C}\right)^{-1}. \qquad \text{(S4)}$$

Equation S4 gives a general construction formula that defines a channel by fixing the parameters $w_u, w_c$, and $L_c$ (equation 16). The expected extension rates result from factor $A$ by setting a flow rate and channel height.

#### Log-logistic growth function fit to Latrunculin B dose response curve

The Young's modulus data shown in figure 5B was well described with a log-logistic growth function as proposed in Urbanska et al:[4]

$$E = E_{\text{lower}} + \frac{E_{\text{upper}} - E_{\text{lower}}}{1 + \exp\left(a\left[\ln\left(\frac{c_{\text{LatB}}}{[\text{nM}]}\right) - \ln\left(\frac{\text{EC}_{50}}{[\text{nM}]}\right)\right]\right)}, \qquad \text{(S5)}$$

with the LatB concentration, $c_{\text{LatB}}$, the lower bound, $E_{\text{lower}}$, and upper bound, $E_{\text{upper}}$, the steepness, $a$, and the effective $\text{EC}_{50}$ dose at which half-maximum response is obtained. The resulting values for all flow rates are given in table S2. The resulting $\text{EC}_{50}$ values are in the range of 10-13 nM and in line with previous reports.[4,5]

#### References


1. Otto, O. *et al.* Real-time deformability cytometry: on-the-fly cell mechanical phenotyping. *Nat Methods* **12**, 199–202 (2015).

2. Wittwer, L. D., Reichel, F., Müller, P., Guck, J. & Aland, S. A new hyperelastic lookup table for RT-DC. *Soft Matter* **19**, 2064–2073 (2023).

3. Büyükurgancı, B. *et al.* Shear rheology of methyl cellulose based solutions for cell mechanical measurements at high shear rates. *Soft Matter* **19**, 1739–1748 (2023).

4. Urbanska, M. *et al.* A comparison of microfluidic methods for high-throughput cell deformability measurements. *Nat Methods* **17**, 587–593 (2020).

5. Gerum, R. *et al.* Viscoelastic properties of suspended cells measured with shear flow deformation cytometry. *Elife* **11**, (2022).

#### Supplementary tables

Table S1: Characterization of PAAm beads. Short names show internal labels for different bead types. Diameters were measured by brightfield microscopy after production. Young's moduli were measured with RT-DC (see Fig. S1).

| Short name | Total monomer concentration [$\%c_T$] | Diameter (mean ± SD) [µm] after production | Young's modulus (mean ± SEM) [Pa] |
|---|---|---|---|
| **S4_12µm** | 4.5 | 12.7 ± 0.7 | 300 ± 2 |
| **S4_17µm** | 4.5 | 17.1 ± 0.6 | 379 ± 17 |
| **S4_18µm** | 5.2 | 18.7 ± 0.6 | 669 ± 32 |
| **S8_15µm** | 5.2 | 14.7 ± 0.8 | 832 ± 15 |
| **S0_3 Set 1** | 6.0 | 13.7 ± 0.4 | 1819 ± 77 |
| **S0_3 Set 2** | 6.0 | 14.5 ± 0.4 | 1846 ± 66 |
| **S0_3 Set 3** | 6.0 | 16.0 ± 0.4 | 1493 ± 17 |
| **S0_3 Set 4** | 6.0 | 16.6 ± 0.4 | 1347 ± 19 |

Table S2: Fit values for the dose response of HL60 cells' Young's moduli as function of LatB concentration (Fig. 5B) fitted with equation S5.

| Flow rate [µL/s] | $\mathbf{EC_{50}}$ [nM] | $\boldsymbol{E}_{\mathbf{lower}}$ [Pa] | $\boldsymbol{E}_{\mathbf{upper}}$ [Pa] | $\boldsymbol{a}$ |
|---|---|---|---|---|
| **0.01** | 13.6 ± 3.3 | 101 ± 10 | 222 ± 7 | 1.8 ± 0.6 |
| **0.02** | 9.7 ± 1.4 | 154 ± 9 | 288 ± 9 | 3.5 ± 2.2 |
| **0.03** | 9.1 ± 1.4 | 222 ± 10 | 365 ± 10 | 3.5 ± 2.2 |

#### Supplementary Figures

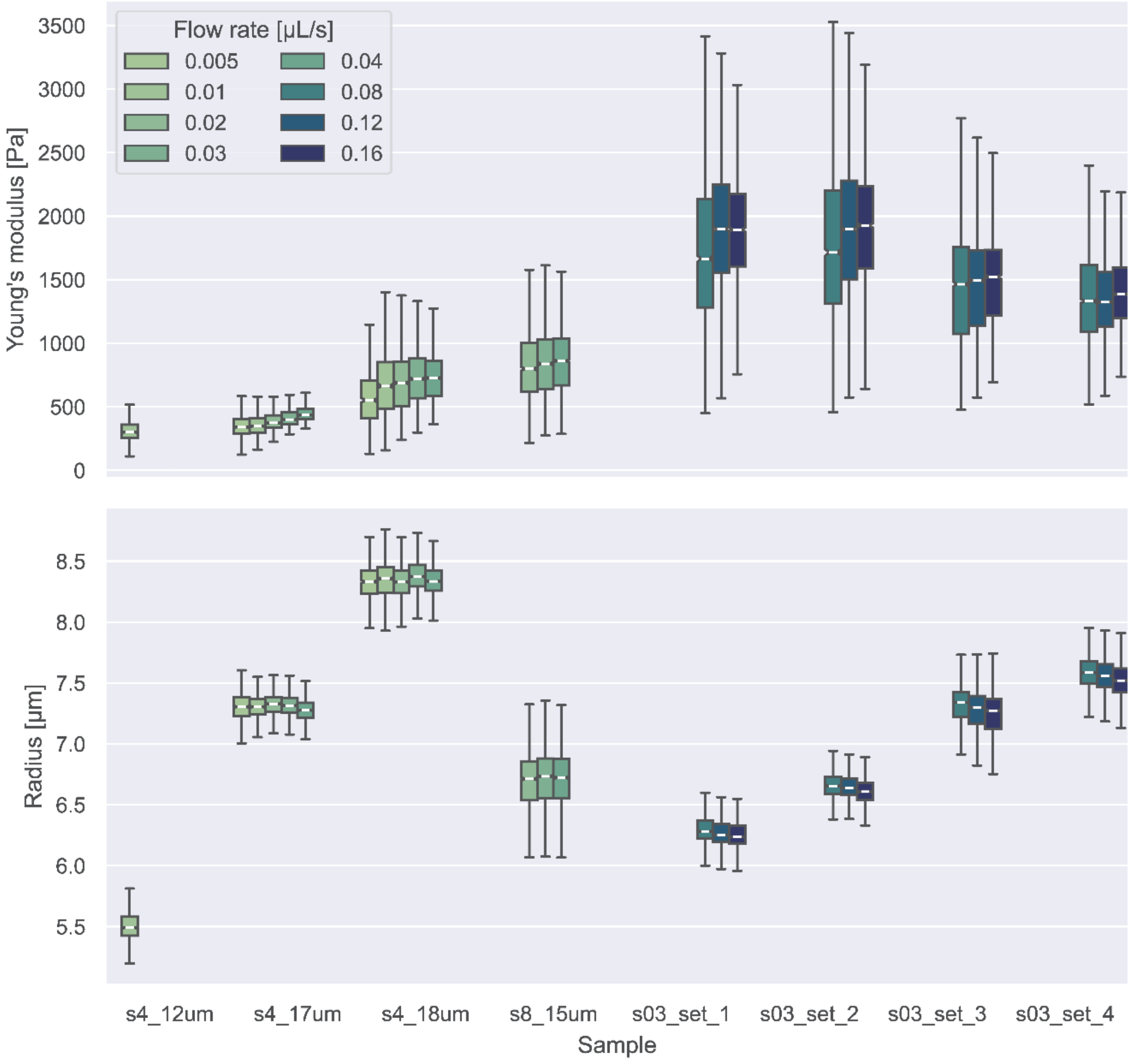


**Figure S1: Young's moduli and radii of PAAm beads from RT-DC measurements**. The radius was calculated from the measured volume, assuming the initial shape was a sphere. Boxes represent the inter-quartile range (IQR). Whiskers show 1.5×IQR borders. Notches around the median indicate the 95% confidence interval. Beads were measured at different flow rates because stiffer beads require higher flow rates to achieve the deformations for accurate computation of the Youngs'modulus.

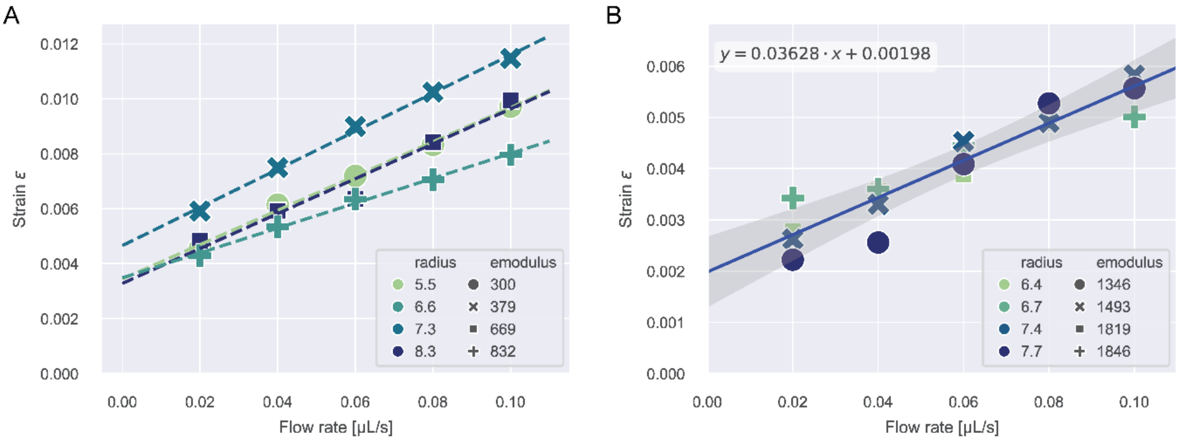


**Figure S2: Strain correction for beads stress measurements** (main text Fig. 2). The strain offset was determined by the intercept of a linear fit to the median strain data vs. flow rate measured in the inlet region. **A**) Strain data and linear fits for beads of type S4 or S8 (see table S1). The resulting intercepts, $\varepsilon_0$, per bead Young's modulus are: $\varepsilon_0(300\ Pa) = 0.0034$, $\varepsilon_0(379\ Pa) = 0.0047$, $\varepsilon_0(669\ Pa) = 0.0033$, $\varepsilon_0(832\ Pa) = 0.0035$ **B**) Strain data and linear fit to the data for beads of type S03. For this bead type, we used the same correction for all samples. The solid line indicates the linear fit to all datapoints and the shaded gray area represents the 2-sigma error band. $\varepsilon_0(S03) = 0.00198$.

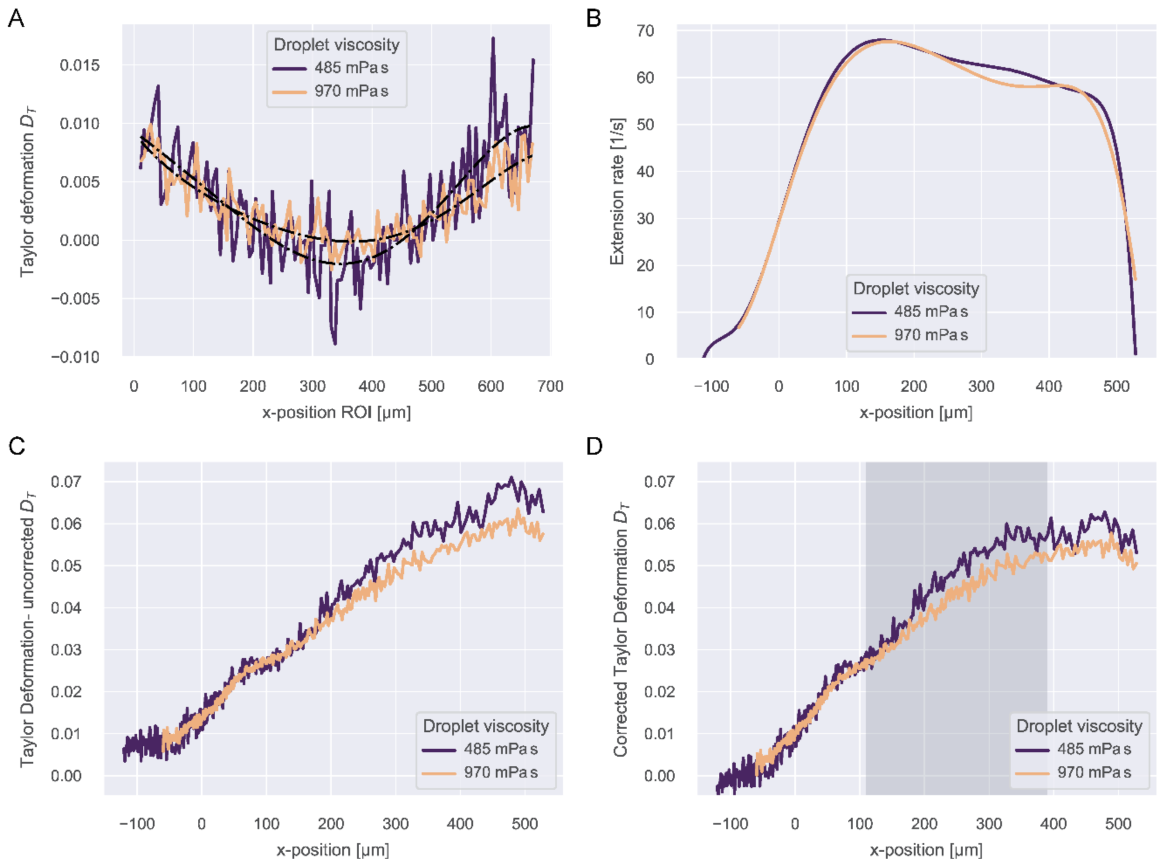


**Figure S3: Strain correction and extension rate for measurements on silicone oil droplets**. **A**) Taylor deformation as function of x in the inlet region of the channel at a flow rate of 0.01 µL/s. The black dash-dotted curves indicate a 6th order polynomial fit to the data. **B**) Extension rate as function of x in the hyperbolic region. **C**) Taylor deformation of the silicone oil droplets before correction. **D**) Corrected Taylor deformation as function of x in the hyperbolic region. The shaded region indicates the analysis region to compute relaxation times, interfacial tension and droplet viscosity. The curve was generated by the data in C), subtracting the values resulting from the polynomial fit shown in A).

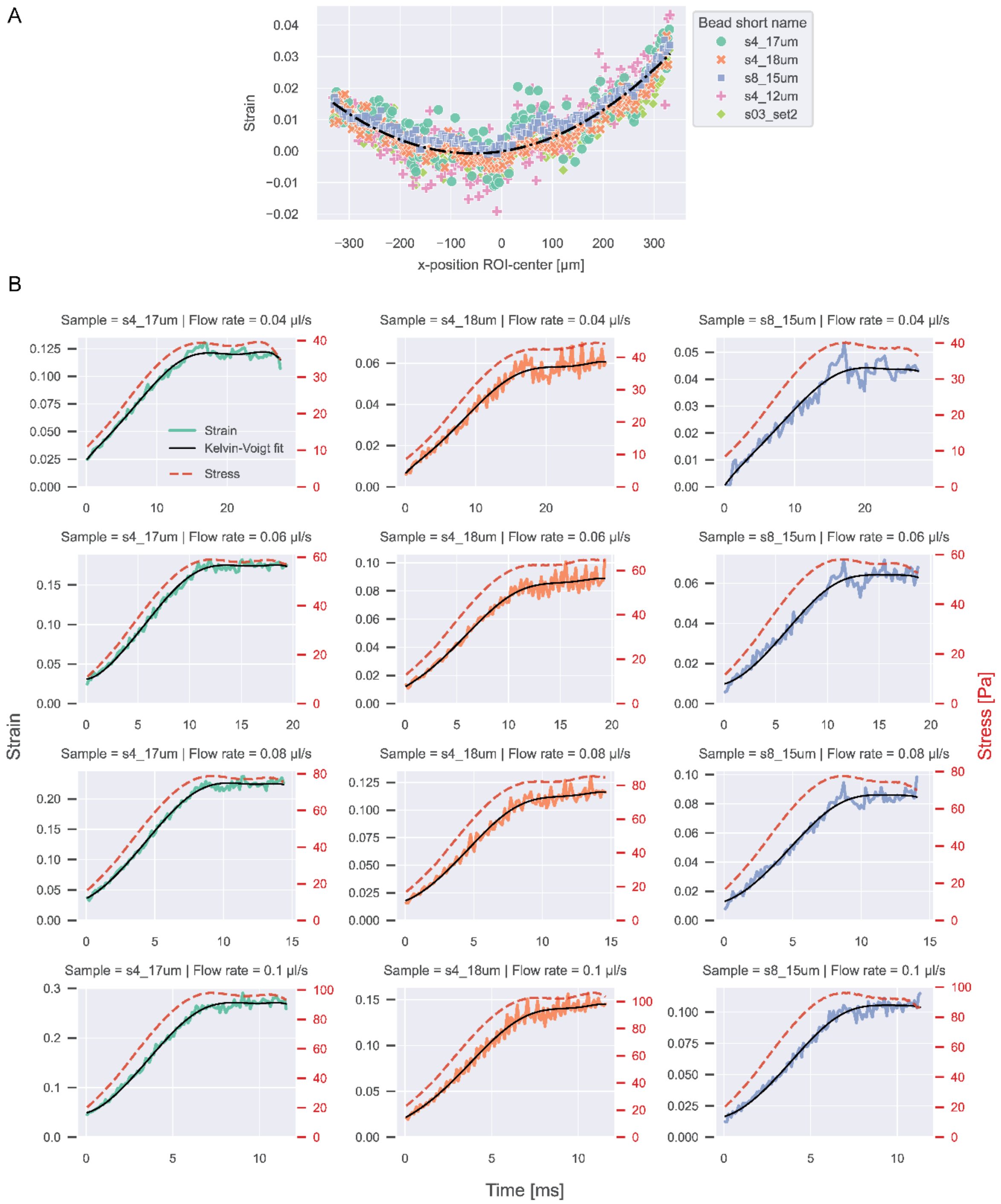


**Figure S4: Strains and stresses of PAAm beads. A**) Strain of different types of PAAm beads in the inlet region of the channel at a flow rate of 0.01 µL/s. We observed that the correction curve was independent of bead type, size, or stiffness. A quadratic function was fitted with x=0 in the center of the ROI: $\varepsilon = 2.1 \cdot 10^{-7}x^2 + 2.4 \cdot 10^{-5}x - 10^{-4}$ (black dash-dotted line). **B**) Corrected strain curves and stress curves as function of time for all bead experiments in the hyperbolic region. The solid black lines indicate the Kelvin-Voigt fit to the data according to equation 13.

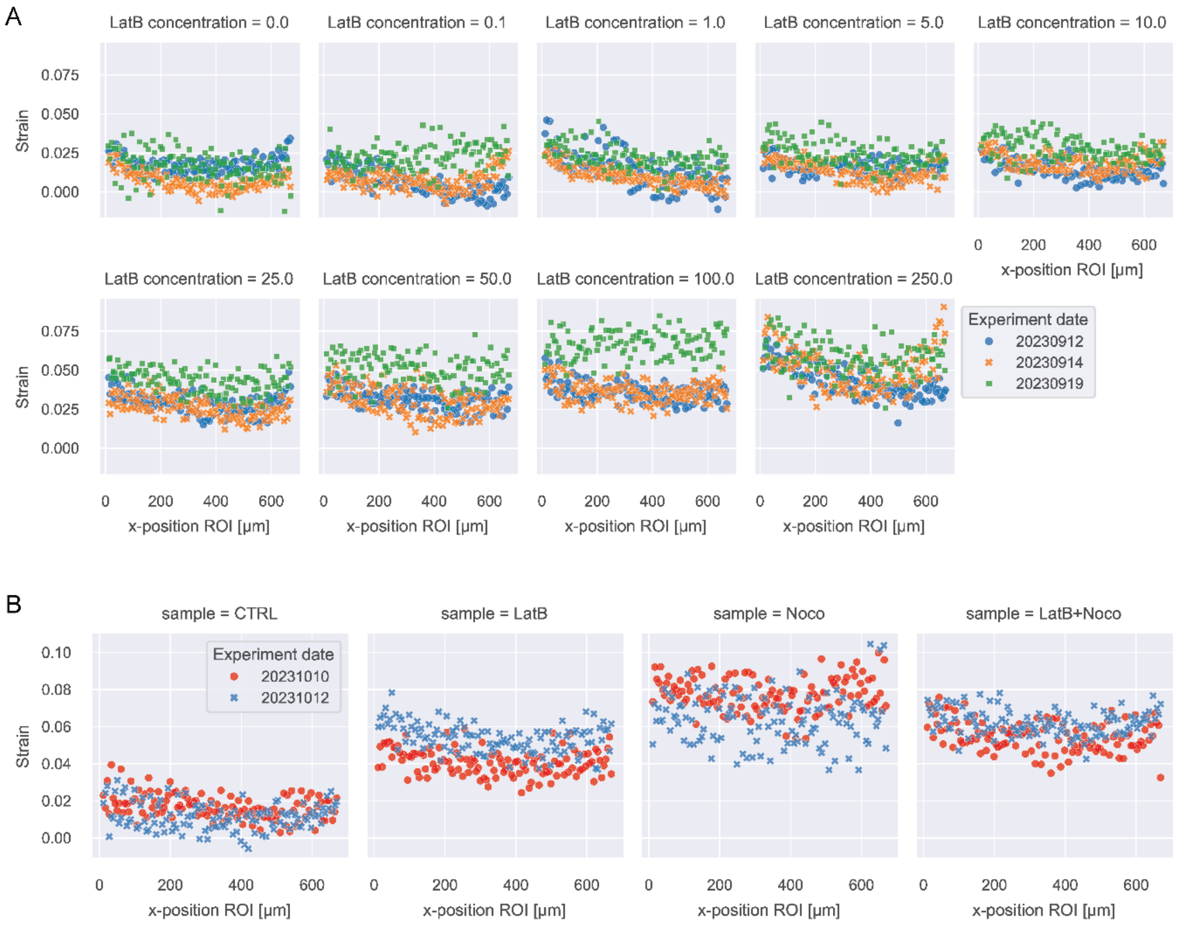


**Figure S5: Strains of HL60 cells + treatment in the inlet region. A**) Strain curves for all HL60 cell with LatB treatment conditions in the inlet region at a flow rate of 0.01 µL/s. **B**) Same data for cells with LatB + Nocodazole treatment.

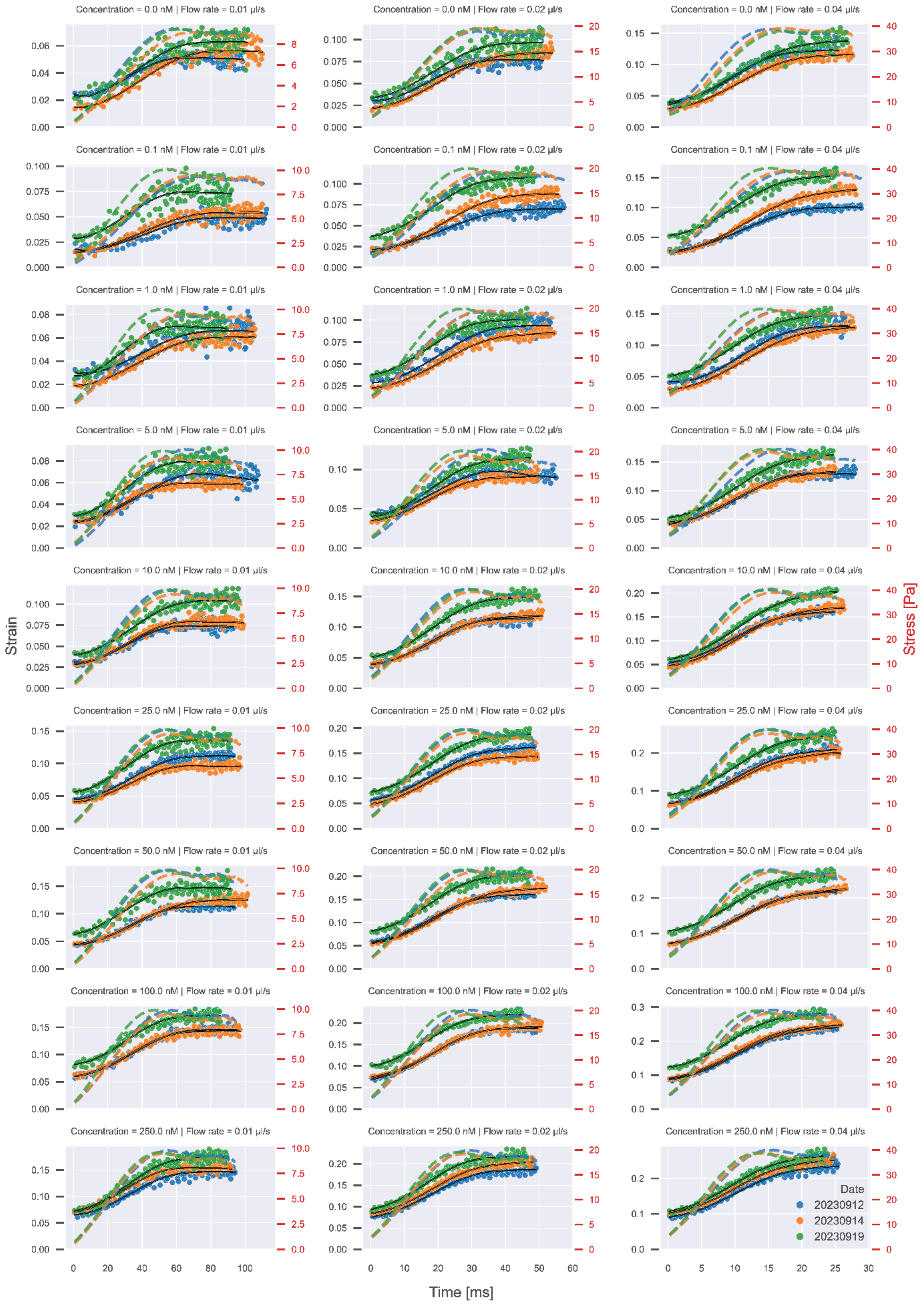


Figure S6: Stress and strain curve as function of time for all HL60 experiments with Latrunculin B treatment. Scatterplots show strain data. Dashed lines show stress curves for the respective experiment. Solid lines show the Kelvin-Voigt fit to the data according to equation 29.

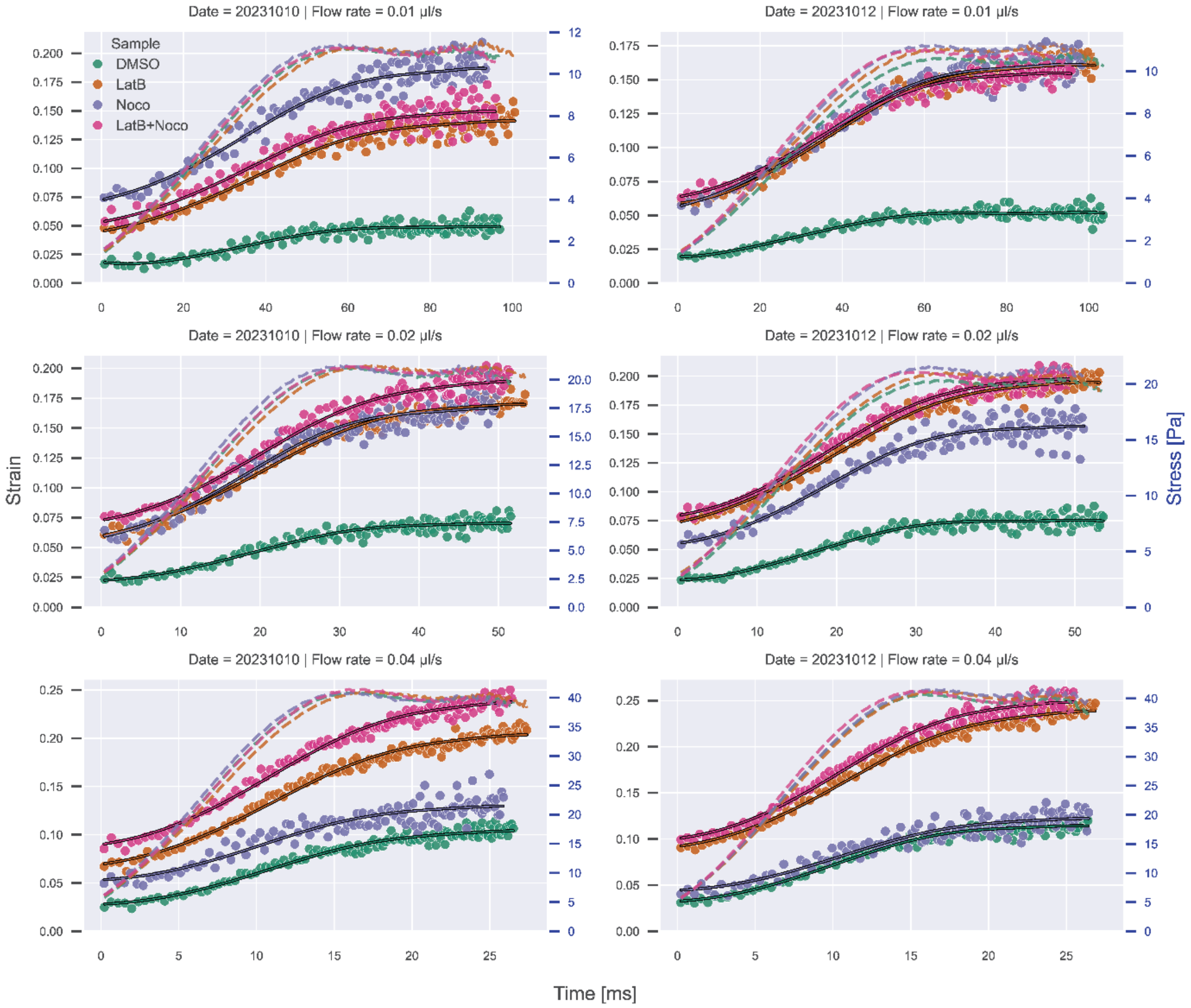


Figure S7: Stress and strain curves as function of time for all HL60 experiments with Latrunculin B and Nocodazole treatment. Scatterplots show strain data. Dashed lines show stress curves for the respective experiment. Solid line show the Kelvin-Voigt fit to the data according to equation 29.
